# Rab34 GTPase mediates ciliary membrane biogenesis in the intracellular ciliogenesis pathway

**DOI:** 10.1101/2020.10.29.360891

**Authors:** Anil Kumar Ganga, Margaret C. Kennedy, Mai E. Oguchi, Shawn D. Gray, Kendall E. Oliver, Tracy A. Knight, Enrique M. De La Cruz, Yuta Homma, Mitsunori Fukuda, David K. Breslow

**Affiliations:** Department of Molecular, Cellular, and Developmental Biology, Yale University, New Haven, CT 06511, USA; Laboratory of Membrane Trafficking Mechanisms, Department of Integrative Life Sciences, Tohoku University, Aobayama, Aoba-ku, Sendai, Miyagi 980-8578, Japan; Department of Molecular Biophysics and Biochemistry, Yale University, New Haven, CT 06511, USA

**Keywords:** cilia, ciliogenesis, ciliopathy, centriole, GTPase, membrane, ciliary vesicle

## Abstract

Primary cilia form by two pathways: an extracellular pathway in which the cilium grows out from the cell surface and an intracellular pathway in which the nascent cilium forms inside the cell. Here we identify the GTPase Rab34 as a selective mediator of intracellular ciliogenesis. We find that Rab34 is required for formation of the ciliary vesicle at the mother centriole and that Rab34 marks the ciliary sheath, a unique sub-domain of assembling intracellular cilia. Rab34 activity is modulated by divergent residues within its GTPase domain, and ciliogenesis requires GTP binding and turnover by Rab34. Because Rab34 is found on assembly intermediates that are unique to intracellular ciliogenesis, we tested its role in the extracellular pathway used by MDCK cells. Consistent with Rab34 acting specifically in the intracellular pathway, MDCK cells ciliate independently of Rab34 and paralog Rab36. Together, these findings reveal a new context-specific molecular requirement for ciliary membrane biogenesis.

## Introduction

The primary cilium is an essential organizing center for signal transduction, and ciliary defects cause congenital disorders known collectively as ciliopathies (Anvarian et al., 2019; Braun and Hildebrandt, 2017; Reiter and Leroux, 2017). Cilia are found on cells in many tissues of the human body, and thus ciliopathy patients exhibit a range of symptoms including retinal degeneration, skeletal defects, kidney cysts, obesity, and intellectual disability (Braun and Hildebrandt, 2017).

Axonemal microtubules form the core of the primary cilium and extend from the mother centriole, which is docked to the plasma membrane in ciliated cells. The axoneme is surrounded by the surface-exposed ciliary membrane, which possesses a distinct protein and lipid composition despite being topologically continuous with the plasma membrane (Garcia et al., 2018; Nachury and Mick, 2019). Although these features of primary cilia are broadly shared, vertebrate cilia can assemble by two distinct pathways that are utilized in a tissue- and cell type-specific manner (Bernabe-Rubio and Alonso, 2017; Breslow and Holland, 2019; Wang and Dynlacht, 2018). These pathways are termed the intracellular pathway and the extracellular pathway and exhibit important differences in the early steps of cilium assembly. As first identified in electron microscopy studies by Sorokin (Sorokin, 1962; Sorokin, 1968), a key initial step in the extracellular pathway is the migration of the mother centriole to the cell periphery, where it docks on the plasma membrane via its distal appendages. The cilium then grows out from the cell surface, driven by the coordinated growth of the axoneme and the surrounding ciliary membrane.

A defining feature of intracellular ciliogenesis is that the nascent cilium forms inside the cytoplasm before becoming exposed to the external environment. Formation of this intracellular intermediate is a complex, step-wise process in which vesicles are first captured at the mother centriole (see Figure 2 A for overview). Subsequent fusion and remodeling of these vesicles yields a ciliary membrane precursor that possesses two contiguous but distinct domains: an axoneme-facing domain that becomes the ciliary membrane and a cytoplasm-facing ciliary sheath (Insinna et al., 2019; Lu et al., 2015; Wu et al., 2018). Ultimately, the ciliary sheath fuses with the plasma membrane, giving rise to a mature surface-exposed cilium. Cilia formed in this manner are commonly recessed within a ciliary pocket that derives from the ciliary sheath. Despite recent progress, several aspects of intracellular ciliogenesis are poorly understood. Specifically, how vesicles captured at the mother centriole give rise to the nascent ciliary membrane is not known. Similarly, how the ciliary membrane and ciliary sheath domains grow and establish their distinct identities is not yet understood. Finally, the relatedness of the intracellular and extracellular pathways is not clear. For example, it is not known if distinct protein machineries are needed for these two modes of ciliogenesis or if similar proteins are involved in both cases but operate in a different order or in different contexts.

Rab family GTPases have widespread roles in intracellular membrane trafficking and organelle biogenesis (Pfeffer, 2017; Stenmark, 2009; Wandinger-Ness and Zerial, 2014). Cilium assembly and ciliary protein trafficking are similarly dependent on Rab proteins, including Rab8, Rab11, and Rab23. These Rab family members each promote ciliary membrane formation and localize to the ciliary membrane or ciliary base (Blacque et al., 2018; Gerondopoulos et al., 2019; Knodler et al., 2010; Leaf and Von Zastrow, 2015; Nachury et al., 2007). Recently, we and others conducted genome-wide CRISPR screens that found *Rab34* to be required for Hedgehog (Hh) signaling (Breslow et al., 2018; Pusapati et al., 2018), which is strictly dependent on cilia in vertebrates (Bangs and Anderson, 2017). Concordantly, *Rab34* mutant mice exhibit ciliopathy features such as polydactyly (Dickinson et al., 2016; Xu et al., 2018), and loss of Rab34 causes ciliary defects (Oguchi et al., 2020; Pusapati et al., 2018; Xu et al., 2018). Despite these recent findings, Rab34 remains one of the least characterized members of the Rab family. It has an unusually divergent sequence, its effectors and regulators are largely unknown, and its precise role at cilia remains elusive.

Here we report that Rab34 promotes ciliogenesis and ciliary membrane formation in the intracellular ciliogenesis pathway. We show that GTP binding and hydrolysis are necessary for Rab34 function and identify Rab34 as the first GTPase that localizes to the ciliary sheath surrounding nascent intracellular cilia. Lastly, MDCK cells that use the extracellular ciliogenesis pathway assemble cilia independently of Rab34, thus revealing Rab34 as a novel context-specific mediator of cilium assembly.

## Results

### Rab34 is required for ciliogenesis

We previously identified *Rab34* as a hit in a genome-wide screen for regulators of cilium-dependent Hh signaling (Breslow et al., 2018). This pooled-format CRISPR screen was based on an NIH-3T3 reporter cell line in which a blastidicin resistance gene is induced in response to Hh pathway activation. Hit genes were then determined by identifying single guide RNAs (sgRNAs) that became depleted upon blasticidin selection relative to non-targeting control sgRNAs. Notably, all ten sgRNAs targeting mouse *Rab34* were strongly depleted upon blasticidin selection, and *Rab34* was a top-scoring hit, similar to core ciliary and Hh pathway genes (Figure 1 A and 1 B).

**Figure 1.**
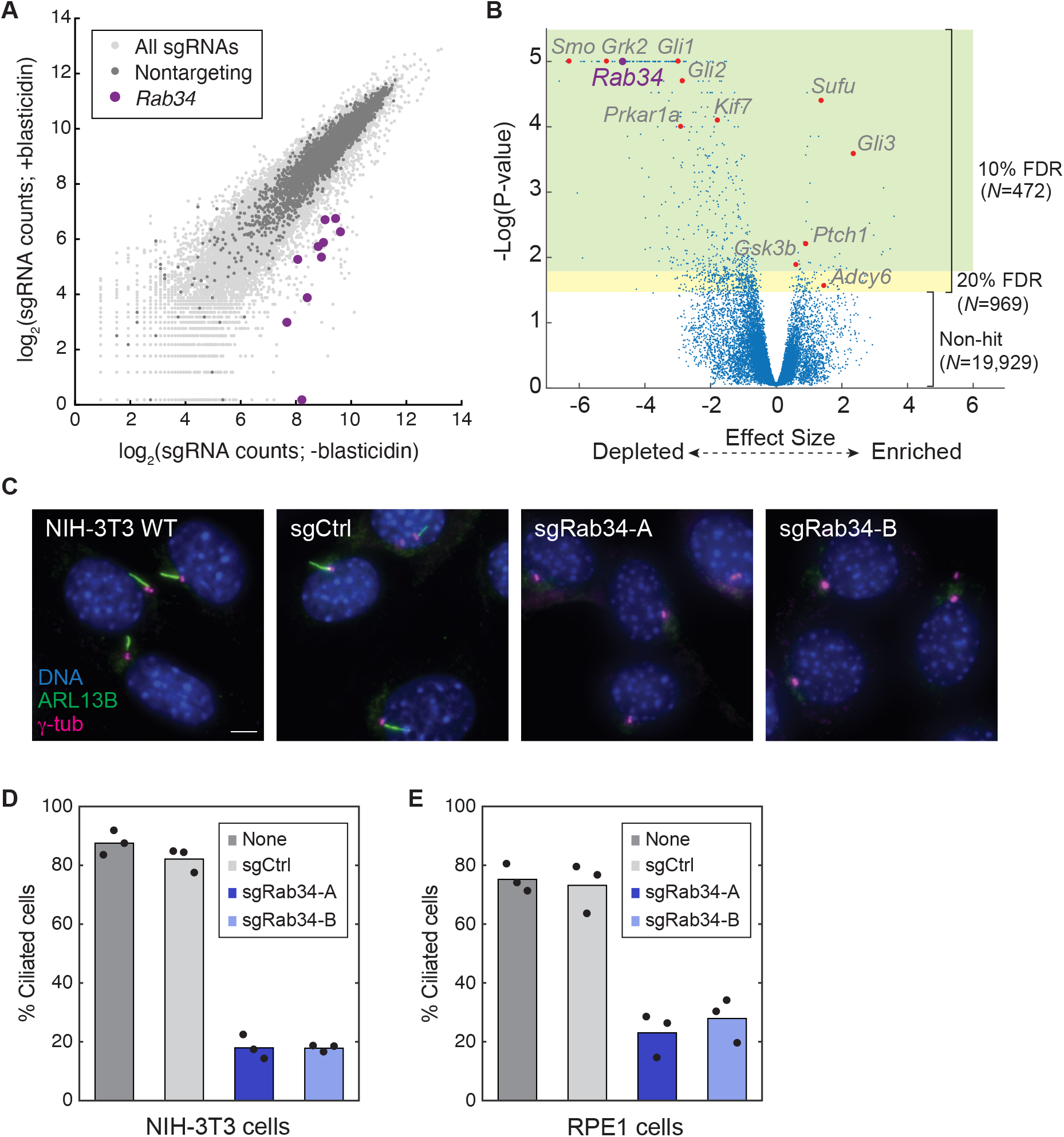
Rab34 GTPase is required for ciliogenesis in NIH-3T3 and RPE1 cells. **A**) Results of a genome-wide CRISPR screen for Hedgehog (Hh) signaling (Breslow et al., 2018) in which signaling-deficient mutants become hypersensitive to blasticidin. NIH-3T3 cells transduced with any of the ten sgRNAs targeting *Rab34* are significantly depleted upon blastidicin treatment compared to non-targeting controls. **B**) Overview of screen results, with all genes plotted according to the confidence that each gene is a hit (-log(P-value)) and the estimated magnitude of the effect on Hh reporter induction and sensitivity to blastidicin (Effect Size). Core Hh pathway components are indicated in red and *Rab34* is indicated in purple. **C**) Cilia (marked by ARL13B) and centrioles (marked by γ-tubulin) were assessed in NIH-3T3 cells transduced with the indicated sgRNAs following 24h serum starvation. **D**) Quantification of ciliogenesis in NIH-3T3 cells described in (**C**); bars indicate mean and dots show values from >120 cells analyzed in each of *N*=3 independent experiments. **E**) Quantification of ciliogenesis in RPE1 cells transduced with the indicated sgRNAs and stained as in (**C**) following 48h serum starvation; bars indicate mean and dots show values from >130 cells analyzed in each of *N*=3 independent experiments.

**Figure 2.**
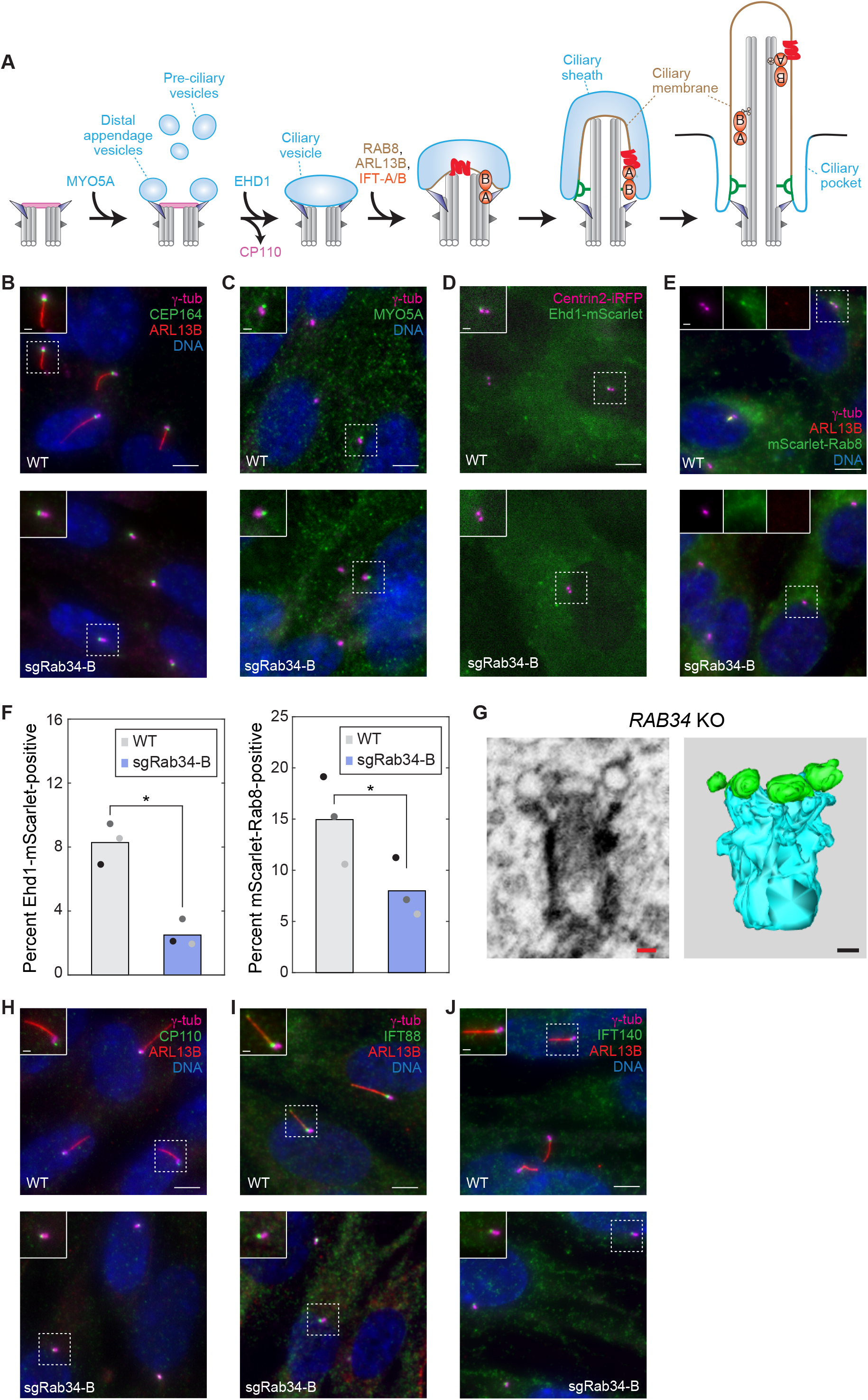
Rab34 is required for formation of the ciliary vesicle from distal appendage vesicles. **A**) Schematic illustration of the intracellular ciliogenesis pathway, highlighting the step-wise formation of a ciliary vesicle associated with the mother centriole’s distal appendages, the elaboration of this membrane and the axonemal microtubules to yield a nascent intracellular cilium, and the subsequent fusion of the ciliary sheath with the plasma membrane that exposes the cilium to the external environment. IFT-complexes are indicated in orange and a ciliary membrane protein in red. **B-E**) The indicated markers were examined in RPE1 WT and *RAB34* sgRNA-B cells. Distal appendage protein CEP164 and pre-ciliary vesicle trafficking motor MYO5A were immunostained after 48 h and 2 h serum starvation, respectively. Cells stably expressing Ehd1-mScarlet and Centrin-miRFP-670 were imaged live 4h after serum starvation, and mScarlet-Rab8 cell lines were stained 24h after serum starvation. Scale bars: 5 μm (insets: 1μm). **F**) The percentage of cells exhibiting localization of Ehd1-mScarlet (left) and mScarlet-Rab8 (right) to the mother centriole or cilium is shown. Bars indicate mean and dots show values *N*=3 independent experiments. P-values determined by two-sided paired t-test are 8.18×10^−3^ and 2.19×10^−2^ for Ehd1 and Rab8, respectively. **G**) Volumetric analysis of centrioles and associated membranes by focused ion beam scanning electron microscopy (FIB-SEM) was conducted in *RAB34* knockout RPE1 cells (sgRab34-B pool). 7×7×7nm voxels were acquired and individual slices as well as 3D reconstructions are shown (centrioles are segmented in cyan and vesicles in green). Scale bars indicate 100nm. See also Supplementary Figure 1 E. **H-J**) Centriolar capping protein CP110, IFT-B component IFT88, and IFT-A component IFT140 were examined after 48h serum starvation in RPE1 WT and *RAB34* sgRNA-B cells. Scale bars: 5 μm (insets: 1μm).

To further investigate the role of *Rab34* in cilium-dependent Hh signaling, we selected two sgRNAs from our screen and generated mutant NIH-3T3 cell pools. In both *Rab34* mutant cell pools, ciliogenesis was strongly reduced compared to wildtype cells or cells transduced with a non-targeting control sgRNA (Figure 1 C-D). This defect was observed when either ARL13B or acetylated tubulin was used as the ciliary marker (Supplementary Figure 1 A) and could be rescued by expression of human *RAB34* (see below and Figure 3 C). We next evaluated the role of *RAB34* in the RPE1 cell line, a widely used model for ciliogenesis. We targeted human *RAB34* with two different sgRNAs and generated clonal and polyclonal cell lines in which we observed frame-shift mutations and/or strongly reduced levels of Rab34 protein (Supplementary Figure 1 D). Similar to *Rab34* mutant NIH-3T3 cells, ciliogenesis was also greatly reduced in all RPE1 *RAB34* mutants (Figure 1 E and Supplementary Figure 1 B-C). These results show that *Rab34* has a critical role in ciliogenesis.

**Figure 3.**
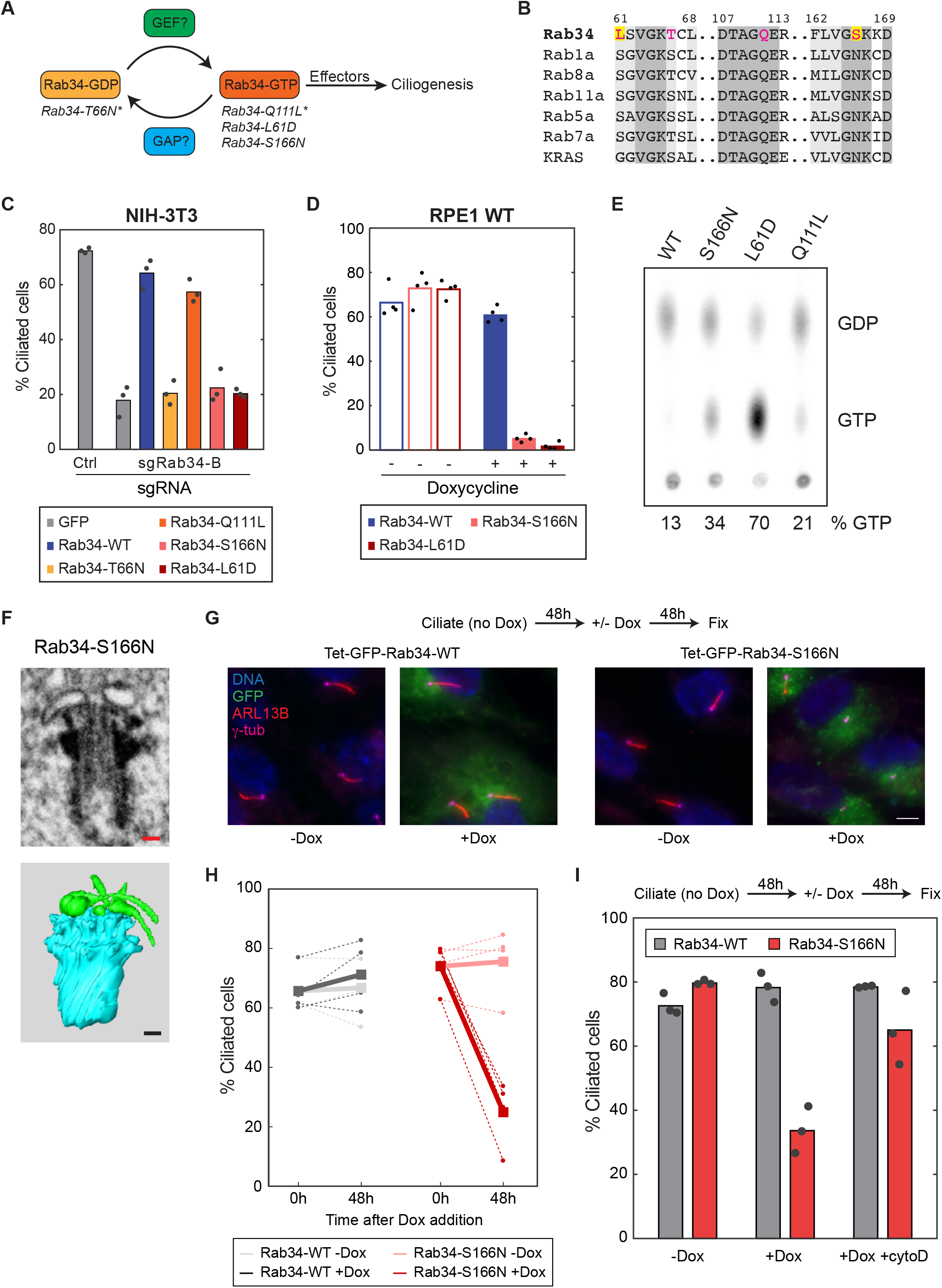
The nucleotide cycle of atypical GTPase Rab34 is required for ciliogenesis and cilium maintenance. **A**) Illustration of the Rab34 nucleotide cycle including Guanine nucleotide Exchange Factors (GEFs), GTPase Activating Proteins (GAPs), and effectors that support its role in ciliogenesis. Mutants of interest and their nucleotide state are shown (predicted nucleotide states are denoted with asterisks; see also panel **E** for observed nucleotide states). **B**) Alignment of key regions of human Rab34 with other human Rab GTPases and KRAS. Conserved residues T66 and Q111 and divergent residues L61 and S166 are indicated. **C**) NIH-3T3 cells transduced with the indicated sgRNAs were transfected with GFP-FKBP or the indicated GFP-Rab34 constructs. Ciliogenesis was assessed as in Figure 1C; bars indicate mean and dots show values from >40 cells analyzed in each of *N*=3 independent experiments. **D**) Ciliogenesis was assessed in RPE1 stable cell lines expressing the indicated GFP-Rab34 variants under control of a doxycycline-inducible promoter. Where indicated, doxycycline was included for 24h preceding serum starvation and during serum starvation. Bars indicate mean and dots show values from *N*=4 independent experiments. **E**) Analysis of GTP and GDP bound to GFP-Rab34 following expression and metabolic labeling in HEK-293T cells, anti-GFP immunoprecipitation, thin layer chromatography of bound nucleotides, and autoradiography. One of 3 representative experiments. **F**) Volumetric analysis of centrioles and associated membranes by focused ion beam scanning electron microscopy (FIB-SEM) was conducted in doxycycline-induced RPE1 cells expressing GFP-Rab34-S166N. 7×7×7nm voxels were acquired as in Figure 2 G. Scale bars indicate 100nm. **G**) RPE1 cell lines with doxycycline-inducible GFP-Rab34 (WT and S166N) were serum-starved for 48h in the absence of doxycycline to allow ciliogenesis to proceed. Cells were then serum starved for an additional 48h in the presence or absence of doxycycline before immunostaining for cilia (ARL13B) and centrioles (γ-tubulin). Scale bars: 5 μm (insets: 1μm). **H**) Quantification of cilia maintenance assay described in (**G**). Dashed lines indicate values from independent experiments and thick, solid lines indicate mean of *N*=3 replicates; dark colors indicate doxycycline-treated cells and light colors indicate untreated cells. **I**) As in (**G**), cells were allowed to ciliate in the absence of doxycycline before 48h of doxycycline treatment in the presence or absence of 100 nM cytochalasin D (cytoD). Ciliogenesis was quantified, with dots indicating values from individual experiments and bars indicating mean from *N*=3 independent experiments.

### Rab34 mediates ciliary vesicle formation

The intracellular ciliogenesis pathway used by NIH-3T3 and RPE1 cells involves a number of sequential steps (Figure 2 A). We therefore asked at what stage in ciliogenesis *RAB34* mutant cells are blocked, focusing first on membrane trafficking and remodeling events given the widespread roles of Rab’s in these processes. One of the earliest events in intracellular ciliogenesis is transport of pre-ciliary vesicles to the mother centriole by MYO5A, where they are captured at the distal appendages to give rise to MYO5A-labeled Distal Appendage Vesicles (DAVs) (Lu et al., 2015; Schmidt et al., 2012; Wu et al., 2018). These events appear to occur normally in *RAB3*4 mutant cells, as distal appendages marked by CEP164 are present, and MYO5A is still recruited to the mother centriole (Figure 2 B-C). After DAVs are captured at the mother centriole, they fuse into a larger ciliary vesicle through the action of EHD family ATPases along with Pascin and SNAP29 (Insinna et al., 2019; Lu et al., 2015). Notably, *RAB34* knockout cells exhibited a significant defect in recruitment of Ehd1-mScarlet-I to the mother centriole (Figure 2 D, F). A similar defect was also observed for the GTPases Rab8 and ARL13B (Figure 2 E-F and Supplementary Figure 1 B), which are normally recruited after EHD1 to promote ciliary membrane growth (Bernabe-Rubio and Alonso, 2017; Nachury and Mick, 2019). To directly observe the membrane remodeling events that occur during intracellular ciliogenesis, we used 3D reconstructions from Focused Ion Beam Scanning Electron Microscopy (FIB-SEM) at an isotropic 7 nm resolution. Strikingly, reconstructions from *RAB34* mutant cells consistently revealed multiple small vesicles docked to the distal appendages without any evident ciliary vesicle (Figure 2 G and Supplementary Figure 1 E). Thus, DAVs are recruited normally in *RAB34* mutant cells, but Rab34 is required for DAVs to fuse and form a ciliary vesicle. This phenotype is similar to that seen upon *EHD1* knockdown (Lu et al., 2015) and is consistent with our observed role for Rab34 in recruiting EHD1 to the mother centriole.

In addition to ciliary membrane formation, ciliogenesis also requires removal of centriolar capping protein CP110 from the mother centriole and recruitment of the intraflagellar transport (IFT) complexes IFT-A and IFT-B (Breslow and Holland, 2019). *RAB34* mutant cells appear to remove CP110 normally and to recruit IFT88 and IFT140, components of the IFT-B subunit and IFT-A complexes, respectively (Figure 2 H-J). Although *EHD1* knockdown was previously reported to impair CP110 removal (Lu et al., 2015), our results indicate that CP110 removal and IFT complex recruitment can occur independently of Rab34-dependent ciliary membrane formation.

### Rab34 GTP binding and hydrolysis are necessary for ciliogenesis and cilium maintenance

Rab GTPases direct vesicular trafficking and membrane remodeling by cycling between an inactive GDP-bound state and an active GTP-bound state that associates with effectors (Figure 3 A). Conserved residues required for GDP binding and GTP hydrolysis have been well characterized in Rab family GTPases and have been exploited to generate mutant alleles that alter the nucleotide cycle in defined ways (Lee et al., 2009; Muller and Goody, 2018). In the case of Rab34, Gln111 of the DXXGQA motif in the switch II region is predicted to be essential for GTPase activity; therefore, a Q111L mutation is predicted to inhibit GTP hydrolysis and lock Rab34 in the GTP-bound state. Conversely, Thr66 of the GxxxxGK(S/T) motif in the P-loop is needed for GTP binding, and a T66N mutation is expected to shift Rab34 to the GDP-bound (or nucleotide-free) state (Figure 3 A-B). To test the role of the Rab34 nucleotide cycle in ciliogenesis, we transfected control and *Rab34* mutant NIH-3T3 cells with GFP-tagged versions of Rab34-WT, Rab34-T66N, and Rab34-Q111L. While Rab34-WT and Rab34-Q111L efficiently rescued the *Rab34* mutant ciliogenesis defect, Rab34-T66N did not, showing that GTP-binding is essential for ciliogenesis (Figure 3 C). Rab34-T66N expression in wildtype cells did not block ciliogenesis, in contrast to the dominant-negative effect commonly observed for such P-loop mutants (data not shown).

While analyzing the sequence of Rab34, we noticed two residues—Leu61 and Ser166— where Rab34’s sequence is conserved across species but diverges from that of other Rab GTPases (Figure 3 B and Supplementary Figure 2 A). Moreover, these residues are likely to be functionally important: Leu61 lies in the P-loop and corresponds to Gly12 of KRAS, where G12D is a common oncogenic mutation (Prior et al., 2012). Ser166 is typically an asparagine in the nucleotide-contacting NKxD motif of small GTPases, and mutations at this position impair nucleotide binding (Tisdale et al., 1992; Walworth et al., 1989). To test the functional significance of these non-canonical residues in Rab34, we examined ciliogenesis in *Rab34* mutant cells expressing Rab34-S166N (reverting Ser166 to the canonical Asn) and GFP-Rab34-L61D (mimicking KRAS-G12D). Strikingly, Rab34-S166N and Rab34-L61D not only failed to rescue ciliogenesis in *Rab34* cells (Figure 3 C) but also appeared to reduce ciliogenesis in wildtype cells (data not shown). To further examine this potential dominant-negative effect, we generated RPE1 cells stably expressing doxycycline-inducible GFP-Rab34-WT, L61D, and S166N. While induced Rab34-WT cells assembled cilia efficiently, Rab34-L61D and Rab34-S166N induction caused a severe ciliogenesis defect (Figure 3 D). Further characterization revealed that Rab34-S166N did not inhibit CEP164 localization, CP110 removal, or IFT complex recruitment at the mother centriole, matching our observations in *RAB34* knockout cells. FIB-SEM reconstructions showed that cells expressing Rab34-S166N also arrested ciliogenesis with a number of DAVs docked to the mother centriole. However, in this case the DAVs display a striking elongated or tubular morphology (Figure 3 F and Supplementary Figure 2 H). This finding strongly suggests that Rab34 directly shapes the DAV membranes and that Rab34-S166N inhibits membrane remodeling at the mother centriole.

Given Rab34’s non-canonical sequence, it is difficult to predict how the S166N and L61D mutations affect the nucleotide cycle of Rab34. We therefore determined the nucleotide state of these Rab34 mutants in cells by metabolic labeling. Specifically, HEK293T cells transfected with wildtype GFP-Rab34 or mutants of interest were labeled with ^32^P orthophosphate before anti-GFP immunoprecipitation and analysis of bound nucleotides by thin-layer chromatography. Consistent with an earlier report (Sun et al., 2003), Rab34-WT protein was mostly GDP-bound. However, Rab34-S166N and Rab34-L61D both bind more GTP, suggesting that these are GTP-locked mutants (Figure 3 E). By contrast, the Q111L mutant canonically predicted to be GTP-locked was similar to wildtype. Increased GTP binding by Rab34-S166N and Rab34-L61D is further supported by immunoprecipitation experiments in which these mutants exhibit increased binding to RILPL1, a putative effector that recognizes the GTP state (Fukuda et al., 2008; Matsui et al., 2012), and decreased binding to Rab-GDI, a binding partner selective for the GDP-bound form of Rab GTPases (Muller and Goody, 2018) (Supplementary Figure 2 B).

To determine how these mutations affect Rab34’s GTPase activity, we measured GTP hydrolysis by purified recombinant Rab34 and mutants. Notably, Rab34-L61D exhibited a ~100-fold reduced k_cat_ than wildtype Rab34 and a ~100-fold tighter K_m_ for GTP. We detected no measurable GTPase activity for Rab34-S166N under our experimental conditions (Supplementary Figure 2 C). Thus, our enzymatic analysis as well as our cellular studies both indicate that Rab34-L61D and Rab34-S166N are shifted toward a GTP-bound state and indicate that GTP binding as well as hydrolysis by Rab34 are required for ciliogenesis.

Lastly, the ability to inducibly express Rab34-S166N allowed us to examine the effect of disrupting Rab34 function after completion of cilium assembly. Interestingly, induction of Rab34-S166N after cilium assembly resulted in progressive loss of cilia (Figure 3 G-H), while uninduced Rab34-S166N cells and induced Rab34-WT cells retained a high level of ciliation. Rab34 is therefore needed to maintain cilia in cultured cells. Moreover, treating cells with the actin poison cytochalasin D (cytoD) rescued this defect. Thus, the ability of cytoD to stabilize cilia (Mirvis et al., 2018) bypasses the need for Rab34 in cilium homeostasis (Figure 3 I).

### Rab34 dynamically localizes to the mother centriole and cilium during ciliogenesis

Because Rab family GTPases commonly localize to the membrane structures they regulate, we examined Rab34 localization in RPE1 cells after serum starvation for 48 h. Notably, staining with two commercially available antibodies revealed colocalization of Rab34 with ciliary markers ARL13B and polyglutamylated tubulin (polyE-tub) in ~10% of ciliated cells (Figure 4 A, D). We also commonly observed a punctum of Rab34 at the mother centriole in unciliated cells, and similar localizations were also observed in NIH-3T3 cells (Figure 4 B). At earlier stages of ciliogenesis (4 h after serum starvation), we found that ~35% of mother centrioles were Rab34-positive while only ~10% were marked by ARL13B, suggesting that Rab34 is recruited to the mother centriole before ARL13B (Figure 4 C). By contrast, as more ARL13B-positive cilia formed at later timepoints, the fraction of cilia with Rab34 decreased (Figure 4 D). Thus, Rab34 may act early during ciliogenesis and be selectively present on nascent cilia versus mature cilia.

**Figure 4.**
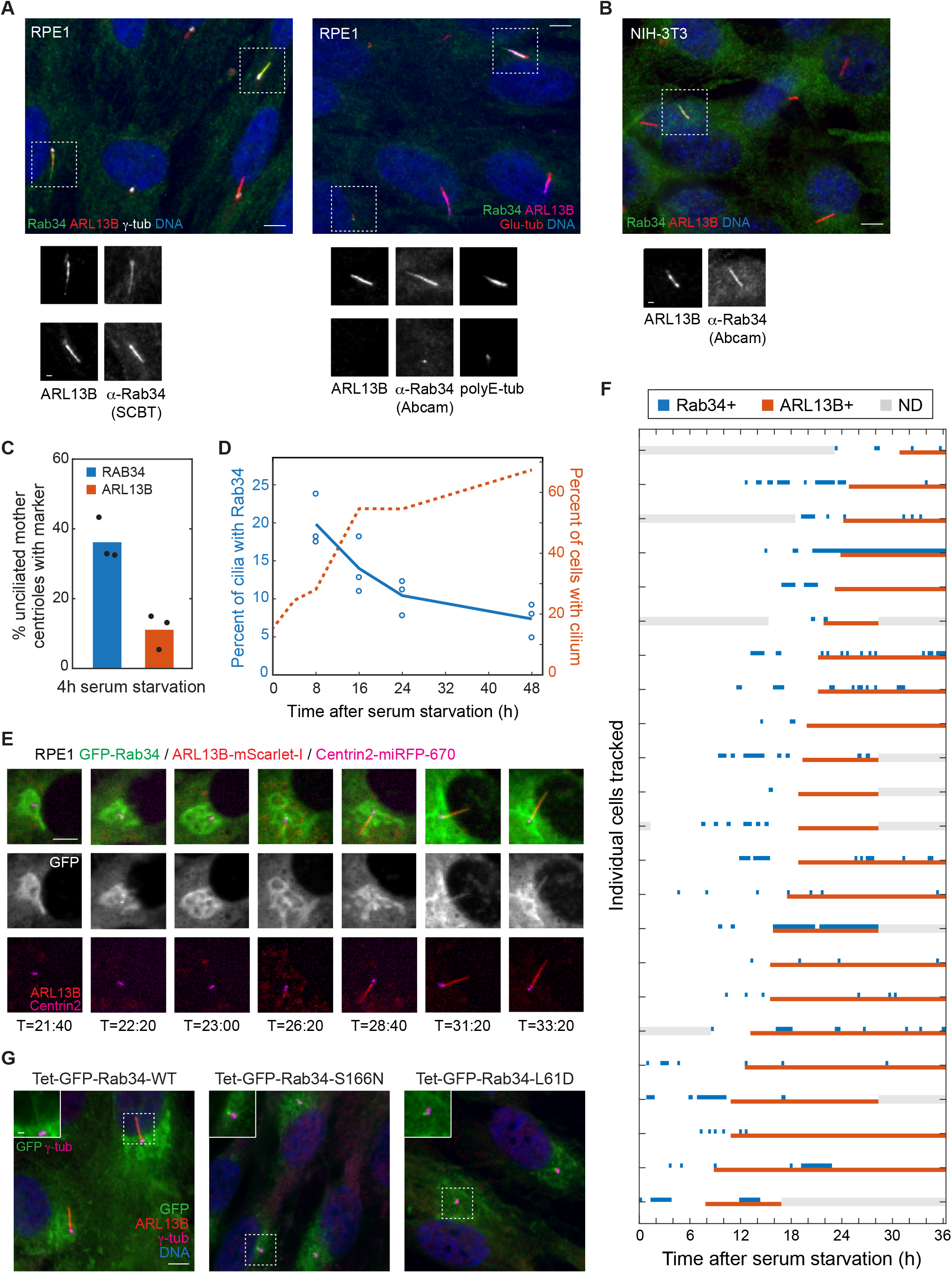
Rab34 dynamically localizes to the mother centriole and cilium during ciliogenesis. **A**) RPE1 cells were stained for the indicated ciliary and centriolar markers as well as for Rab34 using antibodies available from Abcam or Santa Cruz Biotechnology (SCBT). Scale bars: 5 μm (insets: 1μm). **B**) NIH-3T3 cells were stained with the indicated antibodies as in (**A**). **C**) At an early stage of ciliogenesis (4h after serum starvation), the percent of unciliated centrioles that have recruited puncta of Rab34 or ARL13B is plotted. Bars indicate mean and dots show values from each of *N*=3 independent experiments. **D**) The percentage of cilia (marked by ARL13B) that are positive for Rab34 is shown for the indicated times after serum starvation (left-hand Y-axis; the earliest timepoints are omitted due to the low level of ciliation). The fraction of cells with cilia is plotted on the right-hand Y-axis. Circles indicate values from individual experiments and lines show mean from *N*=3 independent experiments. **E**) Time-lapse imaging of RPE1 cell line expressing GFP-Rab34, Arl13b-mScarlet, and centrin2-miRFP-670 was performed, and select images of a representative cell are shown at the indicated times after serum starvation. Scale bar: 5 μm. **F**) Localization of GFP-Rab34 and Arl13b-mScarlet was assessed for 23 ciliating cells tracked for up to 36h after serum starvation. Timepoints when GFP-Rab34 or Arl13b-mScarlet were detected at the mother centriole or cilium are indicated by blue and red boxes, respectively; gray boxes indicate when localization could not be determined (e.g. due to cell leaving field of view); cells are sorted by time of Arl13b arrival and cilium extension. **G**) Localization of GFP-tagged Rab34 was assessed in RPE1 cells stably expressing wildtype Rab34 or the indicated variants. Doxycycline was added prior to and during 48h serum starvation.

To further understand Rab34 localization dynamics, we carried out live-cell imaging in RPE1 cells stably expressing ARL13B-mScarlet-I, Centrin2-miRFP-670, and doxycycline-inducible GFP-Rab34. By monitoring cilium assembly in 23 cells, we were able to reach a number of conclusions (Figure 4 E-F). First, although only a fraction of centrioles and cilia are labeled by Rab34 at any given timepoint, all ciliating cells dynamically recruit Rab34 to the mother centriole during ciliogenesis. Second, Rab34 consistently arrives at the mother centriole several hours before ARL13B. Third, the vast majority of cells exhibit ciliary localization of Rab34 after an ARL13B-positive cilium has formed. Rab34 also appeared to be dynamically gained and lost from cilia, although we note that identifying Rab34-positive cilia was sometimes difficult due to additional GFP-Rab34 localization to the Golgi (in such cases, cilia were scored as Rab34-negative; thus, our analysis likely underestimates Rab34 ciliary localization). Overall, the Rab34 dynamics we observed are consistent with a key role for Rab34 in ciliary membrane biogenesis and with the severe loss of cilia seen in *RAB34* mutant cells.

We also examined the localization of dominant-negative GFP-Rab34-S166N and GFP-Rab34-L61D. Strikingly, these proteins localized as a bright punctum at the mother centriole in ~90% of cells (Figure 4 G and Supplementary Figure 3 A). Given that these cells also exhibit distended ciliary vesicles in our FIB-SEM analysis (Figure 3 F), it is likely that Rab34 is present on DAVs during ciliogenesis and that the dominant-negative GTP-locked mutants become trapped on these structures, impairing DAV fusion.

### Rab34 localization and function are specific to intracellular ciliogenesis

We next investigated why Rab34 localizes to only a subset of cilia and why this fraction declines during the course of ciliogenesis (Figure 4 D). Specifically, we asked whether Rab34 localizes to nascent intracellular cilia. To distinguish intracellular (‘inside’) cilia from extracellular (‘outside’) cilia, we took advantage of the IN/OUT assay (Kukic et al., 2016), in which antibody accessibility is used to assess whether the ciliary membrane protein Smo is exposed to the external environment. Specifically, we used a dye-conjugated anti-GFP Nanobody (Nb) (Kirchhofer et al., 2010) to label Smo tagged on its extracellular N-terminus with pHluorin, a GFP variant, in live cells prior to fixation, permeabilization, and anti-Rab34 immunostaining. Strikingly, even though most (~ 90%) cilia were extracellular after 48h of serum-starvation (Figure 5 B), all of the Rab34-positive cilia were intracellular (Figure 5 A-B). Furthermore, Rab34 localized to the majority of inside cilia (Supplementary Figure 3 B), indicating that Rab34 is a GTPase marker of intracellular cilia. We note that our staining procedure differs from the originally described IN/OUT assay by using the GFP Nb for pHluorin-Smo staining, which we found improved detection of extracellular cilia; however, we obtained similar results with the original protocol, finding that an anti-GFP Ab did not stain 69 out of 70 Rab34-positive cilia (Supplementary Figure 3 C-D). Thus, Rab34 localizes to a small fraction of cilia because it is found exclusively on nascent intracellular cilia.

**Figure 5.**
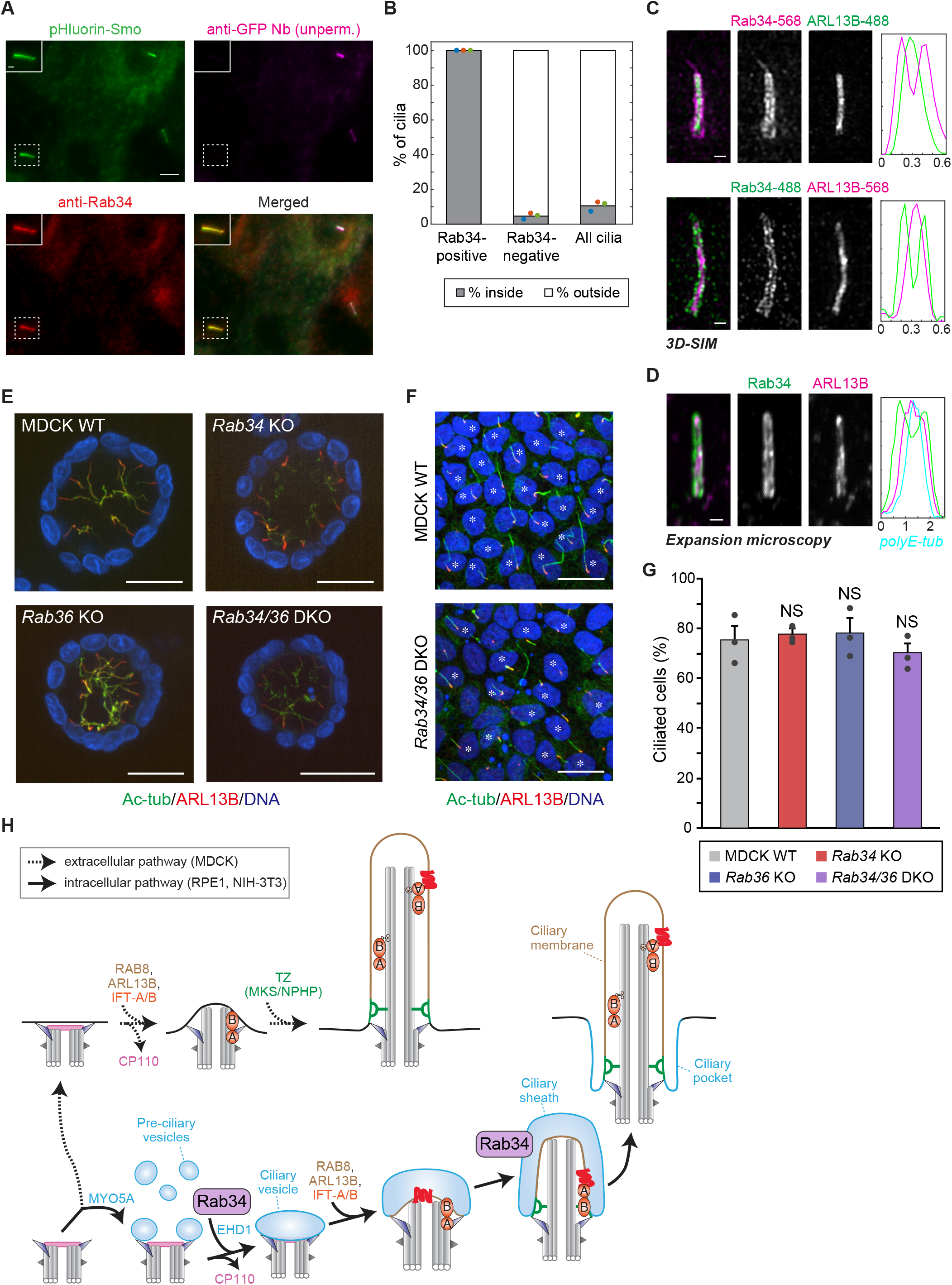
Rab34 localizes to the ciliary sheath of nascent intracellular cilia and is dispensable for the extracellular ciliogenesis used by MDCK cells. **A**) RPE1 cells expressing the pHluorin-Smo In/Out reporter were stained prior to permeabilization with anti-GFP nanobody (Nb) to reveal intracellular versus extracellular cilia followed by permeabilization and staining for Rab34 (Abcam antibody). Inset shows an intracellular (inside) cilium that is positive for Rab34. Scale bars: 5 μm (insets: 1μm). **B**) Quantification of the fraction of cilia that are inside versus outside is shown for the cilia classes indicated, as assessed using the In/Out assay shown in (**A**). Dots indicate values from individual experiments and bars indicate mean from *N*=3 independent experiments encompassing 110 Rab34-positive cilia and 1,651 Rab34-negative cilia. **C**) 3D-SIM (structured illumination microscopy) was performed on RPE1 cells stained for Rab34 and ARL13B. Alexa Fluor 488- and Alexa Fluor 568-conjugated secondary antibodies were used for both target proteins to ensure consistent results independent of emission wavelength. Graphs at right show normalized fluorescence intensity along line-profiles perpendicular to the longitudinal axis of the cilium. Scale bars indicate 100nm. See also Supplementary Figure 3 E. **D**) Expansion microscopy imaging of cells stained for Rab34 and ARL13B. Graph at right shows normalized fluorescence intensity along line-profile perpendicular to the longitudinal axis of the cilium, including for axonemal polyE-tubulin. Scale bar indicates 2μm (after ~4-fold expansion). See also Supplementary Figure 3 F. **E**) Representative images of 3D cysts of parental, *Rab34* KO, *Rab36* KO, and *Rab34*/*36* double-knockout (DKO) MDCK cells. The cells were fixed after 7-day culture in collagen gel and then stained for acetylated tubulin (Ac-tub; green), ARL13B (red), and DAPI (blue). Scale bars, 20 μm. **F**) Representative images of 2D monolayers of parental, *Rab34* KO, *Rab36* KO, and *Rab34*/*36* DKO MDCK cells. The cells were fixed five days after seeding and then stained as in (**E**). Scale bars, 20 μm. Asterisks indicate ciliated cells. **G**) The percentage of ciliated cells in 2D monolayers of parental, *Rab34* KO, *Rab36* KO, and *Rab34*/*36* DKO MDCK cells. Dots indicate values from individual experiments, bars indicate mean from *N*=3 independent experiments, and error bars indicate standard error. Differences that are not significantly different are marked “NS” (Tukey-Kramer test; P > 0.60). **H**) Model for Rab34 function, in which it mediates ciliary membrane formation in the intracellular ciliogenesis pathway (bottom, solid arrows) but not the extracellular ciliogenesis pathway (top, dashed arrows). Rab34 is present on DAVs and the ciliary sheath (blue) but not the ciliary membrane (tan).

To determine whether the ciliary membrane or ciliary sheath harbors Rab34 (Figures 2 A and 5 H), we used 3D structured illumination microscopy (3D-SIM) to resolve these closely apposed membranes. Notably, Rab34 fluorescence consistently surrounds the ARL13B-labeled ciliary membrane (Figure 5 C and Supplementary Figure 3 E). Line scans across the width of the cilium confirmed two peaks of Rab34 surrounding a single ARL13B peak, as seen for other ciliary sheath proteins such as EHD1 and MYO5A (Wu et al., 2018).

To corroborate this finding, we used expansion microscopy, in which fixed samples are embedded in a swellable polymer to allow ~4-fold isotropic expansion after immunostaining, thereby providing an effective 4-fold increase in spatial resolution (Tillberg and Chen, 2019). Using an expansion microscopy protocol optimized for cilia and centrioles (Sahabandu et al., 2019), we again found that Rab34 fluorescence surrounds that of Arl13b, with polyE-tubulin staining of the axoneme marking the central-most part of the cilium (Figure 5 D and Supplementary Figure 3 F). These results establish that Rab34 localizes specifically to the ciliary sheath and identify Rab34 as the first GTPase that is enriched on the ciliary sheath membrane.

As Rab34 localizes to structures and assembly intermediates unique to intracellular ciliogenesis, we hypothesized that Rab34 might be essential for the intracellular pathway but dispensable for the extracellular pathway. To test this possibility, we used CRISPR to knock out *Rab34* in Madin-Darby Canine Kidney II (MDCK) cells, which form polarized epithelia and use the extracellular pathway for ciliogenesis (Bernabe-Rubio et al., 2016; Ghossoub et al., 2011). Consistent with our hypothesis, immunostaining of ciliary markers did not reveal any ciliogenesis defect in *Rab34* mutant MDCK cells grown in either 3D cysts or 2D monolayers (Figure 5 E-F). As Rab34 has high sequence similarity to Rab36 (Supplementary Figure 2 A), we also investigated whether Rab36 might compensate for the loss of Rab34 in MDCK cells. However, like *Rab34* knockout MDCK cells, *Rab36* knockout and *Rab34/Rab36* double-knockout MDCK cells assembled cilia at levels comparable to wildtype control cells (Figure 5E-G). Thus, extracellular ciliogenesis in MDCK cells is independent of both Rab34 and Rab36, while RPE1 and NIH-3T3 cells require Rab34 for intracellular ciliogenesis. Together, our results show a pathway-specific function for Rab34 in ciliogenesis (Figure 5 H).

## Discussion

Here, we found that Rab34 is a critical mediator of ciliary membrane biogenesis specifically in the intracellular ciliogenesis pathway. Through detailed analysis of *RAB34* knockout and dominant-negative mutants, we establish that Rab34 is needed for DAVs at the mother centriole to fuse and form a larger ciliary vesicle. This finding is supported by our observation that factors acting upstream of this step are recruited normally in *RAB34* mutant cells, while proteins that participate in DAV fusion (e.g. EHD1) or downstream ciliary membrane growth (Rab8 and ARL13B) are not (Figure 2). We note that while our work was in progress, Rab34 was suggested to promote early stages of ciliary membrane formation, but the precise step mediated by Rab34 was unclear (Oguchi et al., 2020; Xu et al., 2018). A key advance in our study comes from the application of volumetric high-resolution FIB-SEM to reconstruct the membrane structures that form (and fail to form) in *RAB34* mutant cells, allowing us to unambiguously determine that cells were arrested at the DAV stage (Figure 2 G). A direct role for Rab34 in shaping and remodeling DAVs is further supported by our observation that DAV morphology is altered in cells expressing dominant-negative Rab34-S166N (Figure 3 F).

Our localization studies provide further evidence that Rab34 mediates ciliary membrane formation. Specifically, imaging of fixed and live cells showed dynamic recruitment of Rab34 to the distal end of the mother centriole before ARL13B is present (Figure 4 C-F). The persistent localization of Rab34-S166N at this site further suggests that Rab34 is present on DAVs (Figure 4 G). Our studies also extend recent reports that Rab34 can sometimes localize to cilia (Oguchi et al., 2020; Xu et al., 2018) by defining the precise subset and sub-domains of cilia where Rab34 is found (Figure 5 A-D). Specifically, the In/Out Assay, 3D-SIM, and expansion microscopy showed that Rab34 is present on the ciliary sheath of nascent intracellular cilia. Given the roles of Rab GTPases as markers of membrane identity, Rab34 may establish and maintain the identity of DAVs and the ciliary sheath membrane, helping to distinguish them from the ciliary membrane and other cellular compartments. Furthermore, because the ciliary sheath is the part of the intracellular cilium that faces the cytoplasm, Rab34 on the ciliary sheath may be critical for vesicular trafficking to the nascent cilium, a process that is likely necessary to fuel membrane growth as the intracellular cilium elongates. Alternatively or in addition, Rab34 may mediate the final fusion of the ciliary sheath with the plasma membrane to expose the cilium to the external environment, a process that remains poorly understood. A clearer picture of Rab34’s roles beyond DAV fusion awaits the development of new tools to inactivate Rab34 rapidly during ciliogenesis. It is also noteworthy that, unlike other proteins that localize to the ciliary sheath, such as EHD1 (Lu et al., 2015), Rab34 appears not to be retained on the ciliary pocket of mature cilia. How Rab34’s localization is dynamically regulated will be a key area for future study.

Our studies also provide insight into the biochemical properties of Rab34, an atypical GTPase with key residues that differ from other Rab family members. We find that these divergent residues have critical roles in the Rab34 nucleotide cycle and identify Rab34-S166N and Rab34-L61D as dominant-negative mutants that impair ciliogenesis (Figure 3 D). These mutants preferentially bind GTP in cells and hydrolyze GTP more slowly than wildtype *in vitro*, indicating that GTP hydrolysis as well as GTP binding are necessary for Rab34 function (Figure 3 E and Supplementary Figure 2 C). On the other hand, the putative GTP-locked mutant Q111L did not appear to bind more GTP or to impair ciliogenesis. As reported for some other Rab GTPases (Langemeyer et al., 2014; Nottingham and Pfeffer, 2014), it is possible that this mutant remains susceptible to GAPs that stimulate GTP hydrolysis. Further insights into the Rab34 enzymatic cycle await the identification of GEFs and GAPs that regulate Rab34. Additionally, crystal structures may shed light on how Rab34’s structure and function are impacted by its unusual sequence and by mutations such as S166N and L61D.

Another future challenge will be identifying the effectors that Rab34 regulates to mediate ciliogenesis. We tested Rilpl1 and Rilpl2, as these proteins associate with GTP-bound Rab34 (Supplementary Figure 2 B) and have been previously linked to cilia (Schaub and Stearns, 2013). However, we found that Rilpl1/2 double-knockout cells neither exhibit ciliogenesis defects nor modulate the *Rab34* mutant phenotype (data not shown). Additionally, Rab34 variants that uncouple ciliogenesis from RILP binding have recently been reported (Oguchi et al., 2020). EHD1 is another plausible candidate effector, though we did not detect co-immunoprecipitation of EHD1 with Rab34 (data not shown). Lastly, an appealing possibility is that Rab34 acts in sequence with another Rab family member to promote ciliogenesis. Such Rab cascades have been reported for other membrane trafficking processes (Mizuno-Yamasaki et al., 2012), and a number of other Rab proteins participate in ciliogenesis (Blacque et al., 2018). We anticipate that our newly identified GTP-locked alleles of Rab34 will be valuable tools to uncover Rab34 effectors and regulators.

Our data indicate that Rab34 acts selectively in the intracellular ciliogenesis pathway and that cells using the extracellular pathway can form cilia independently of *Rab34* and paralogous *Rab36* (Figure 5 E-H). It remains unclear if such pathway selectivity is unique to Rab34 or if other proteins such as EHD1 and MYO5A share this property. A pathway-specific role for Rab34 is consistent with the fact that, unlike many genes needed for ciliogenesis, *Rab34* is not conserved in all ciliated organisms but is restricted to metazoans (Diekmann et al., 2011). A more ancestral Rab may fulfill Rab34’s role in other species, but an intriguing possibility is that Rab34 is selectively present in organisms that employ the intracellular ciliogenesis pathway.

Lastly, our findings provide a framework for understanding the physiologic roles of Rab34 and the phenotypic consequences of its inactivation. Homozygous *Rab34* mutant mice die perinatally and exhibit canonical ciliopathy features such as polydactyly and craniofacial malformations (Dickinson et al., 2016; Xu et al., 2018). However, these phenotypes are notably milder than the early embryonic lethality seen for mutants that cause universal cilium loss (Huangfu et al., 2003); instead, these defects are consistent with disruption of only a subset of ciliated tissues. In the case of the reported loss of cilia in the limb bud (Xu et al., 2018), it is noteworthy that cells of the limb bud mesenchyme have a pronounced ciliary pocket and likely ciliate via the intracellular pathway (Haycraft et al., 2005). In the limb bud and cerebellum, *Rab34* is also a target gene induced by Hh signaling (Lee et al., 2010; Vokes et al., 2007). Although the functional significance of *Rab34* induction is unclear, one possibility is that Rab34 promotes cilium re-formation after mitogenic Hh signaling induces cell division and cilium disassembly. Such a model has recently been proposed for Atoh1, which promotes ciliogenesis in cerebellar granule neuron progenitors and is up-regulated by Hh signaling (Chang et al., 2019). In this way Rab34 may ensure cilium homeostasis, a possibility that is consistent with the disruption of cilium maintenance we observed upon Rab34-S166N expression in cultured cells. Finally, while patients with *RAB34* mutations have not yet been reported, understanding the clinical manifestations of *RAB34* disruption is likely to provide valuable new insights into tissue-specific modes of cilium assembly and into the phenotypic variability seen in ciliopathies.

## Materials and methods

### Cell Culture

NIH-3T3, RPE1-hTERT (RPE1), and HEK-293T cell lines were obtained from ATCC; MDCK-II cells were obtained from Katsuhiko Mikoshiba (ShanghaiTech University) and were maintained in a humidified 37°C incubator with 5% CO_2_. NIH-3T3, HEK-293T, and MDCK-II cells were cultured in DMEM, high glucose (Gibco) supplemented with 10% fetal bovine serum (FBS; Sigma Aldrich), 100 units/ml Penicillin, 100 μg/ml Streptomycin, 2 mM Glutamine, and 1 mM sodium pyruvate (Gibco). RPE-1 cells were cultured in DMEM/F-12 medium (Gibco) supplemented with 10% FBS, 100 units/ml of Penicillin, 100 μg/ml Streptomycin, and 2 mM Glutamine. To induce ciliogenesis, NIH-3T3 cells were starved in 0.5% FBS-containing medium for 24 h, and RPE-1 cells were starved in 0.2% FBS-containing medium for 48 h. For analysis of ciliogenesis of MDCK cells grown in two-dimensional (2D) monolayers, cells were cultured until confluence was achieved (5 days after seeding). Three-dimensional (3D) MDCK cysts were formed by culturing in collagen I gel (KOKEN #IPC-50) as described previously (Homma et al., 2019). Where specified, cells were treated with 100 nM cytochalasin D (Sigma Aldrich) or with 1 μg/ml doxycycline (Fisher Scientific).

### DNA cloning

Gibson assembly, Gateway cloning, and standard molecular biology were used for DNA cloning. Human *RAB34* cDNA was obtained from GE Healthcare (MHS6278-202807876) and cloned into Gateway pENTR plasmid pDONR221 (Invitrogen); *RAB34* mutants were prepared by site-directed mutagenesis of pENTR-Rab34. Plasmids for tetracycline-inducible expression of LAP-Rab34 (LAP consists of GFP, TEV protease site, and S-tag) were constructed by modification of pCW-Cas9 (Addgene #50661, gift from Eric Lander and David Sabatini). Stable expression of fluorescently tagged ciliary and centriolar proteins was achieved by modification of pCW-Cas9 using cDNAs encoding ARL13B (Addgene #40879, gift from Tamara Caspary), EHD1 (gift from Chris Westlake), Rab8 (gift from Maxence Nachury), and Centrin2 (gift from Tim Stearns), miRFP670 (Addgene #79987, gift from Vladislav Verkhusha), and mScarlet-I (Addgene #85044, gift from Dorus Gadella). Plasmids for CRISPR-based knockout were constructed using pMCB320 (Addgene #89359, gift from Michael Bassik) and pMJ179 (Addgene #89556, gift from Jonathan Weissman); see Supplementary Table 2 for sgRNA sequences. Plasmids for transient transfection of *RAB34* were made by Gateway cloning using pEF5-FRT-LAP-DEST (Breslow et al., 2013; Ye et al., 2013); for bacterial protein expression, *RAB34* variants were cloned into pGEX-6P1 (GE Healthcare).

### Lentivirus production and cell line generation

VSVG-pseudotyped lentiviral particles were produced by transfection of HEK293T cells with a lentiviral vector and packaging plasmids (pMD2.G, pRSV-Rev, pMDLg/RRE for sgRNAs; pCMV-ΔR-8.91 and pCMV-VSVG for protein-coding constructs). Following transfection using polyethyleneimine (Polysciences #24765-1), virus-containing supernatant was collected 48 h later and filtered through a 0.45 μm polyethersulfone filter (VWR 28145-505). For protein-coding constructs, lentiviral particles were concentrated 10-fold using Lenti-X Concentrator (Takara Biosystems 631231).

Cells were transduced by addition of viral supernatants diluted to an appropriate titer in growth medium containing 4 mg/ml polybrene (Sigma Aldrich H9268). Following 24 h incubation at 37°C, virus-containing medium was removed; after an additional 24 h, cells were passaged and, where appropriate, selection for transduced cells was commenced by addition of 2.0 μg/ml puromycin or 300 μg/ml G418 (InvivoGen) or 8.0 μg/ml blasticidin (InvivoGen). Alternatively, transduced cells were isolated by FACS using a FACSAria III sorter (Becton Dickinson). Unless otherwise indicated, polyclonal pools of transduced cells were used in subsequent experiments.

For CRISPR-based mutagenesis of NIH-3T3 cells, a cell line stably expressing Cas9 was used (3T3-[Shh-BlastR];Cas9, Breslow et al., 2018). For mutagenesis of RPE1 cells, a cell line stably expressing Cas9 was generated by infection with lentiCas9-Blast (Addgene 52962, gift from Feng Zhang) and selection with blasticidin. Cas9-expressing cells were transduced with sgRNA constructs followed by antibiotic selection or FACS-based sorting of clones. Mutant alleles in knockout clones were assessed by sequencing of genomic DNA extracted using the QIAamp DNA Mini kit (Qiagen). The target locus was amplified using flanking primers (see Supplementary Table 2) and subjected to Sanger sequencing. *Rab34*, *Rab36* and *Rab34*/*Rab36* knockout MDCK cells were described previously (Homma et al., 2019).

### Staining for immunofluorescence microscopy

Staining for immunofluorescence microscopy was performed as described previously (Breslow et al., 2018). Briefly, cells were seeded on acid-washed 12-mm #1.5 coverslips (Fisher Scientific). Cells were either fixed in 4% paraformaldehyde (Electron Microscopy Sciences) for 10 min or −20°C methanol for 10 min or 4% paraformaldehyde for 10 min followed by methanol or 10% TCA for 10 min (for MDCK cysts). Coverslips were permeabilized for 10 min in PBS containing 0.1 % Triton X-100, washed with PBS, and blocked for 20 min in PBS supplemented with 5% normal donkey serum (Jackson Immunoresearch) and 3% Bovine Serum Albumin (BSA). Coverslips were then incubated with appropriate primary antibodies diluted in PBS with 3% BSA at room temperature for 1 h, washed five times with PBS containing 3% BSA, incubated with secondary antibodies for 30-60 min at room temperature, washed again, stained with Hoechst 33258 dye, and mounted on glass slides in Fluoromount-G mounting medium (Electron Microscopy Sciences). MDCK cysts were transferred in PBS to glass-bottom dishes for imaging. Primary and secondary antibodies used are listed in Supplementary Table 1.

### In/Out Assay

In/Out Assay with anti-GFP antibody was carried out as described previously (Kukic et al., 2016). Briefly, coverslips were fixed in 4% paraformaldehyde for 10 min and blocked in PBS containing 5% normal donkey serum and 3% BSA for 20 min. After blocking, unpermeabilized coverslips were incubated with rabbit anti-GFP antibody for 30 min at room temperature. After washing, coverslips were fixed again in 4% paraformaldehyde, permeabilized with PBS containing 0.1% Triton X-100, blocked again, and incubated with anti-Rab34 antibody (SCBT) for 1 h at room temperature. Coverslips were then washed, stained with secondary antibodies, and mounted on glass slides as described above.

The In/Out Assay with anti-GFP nanobody (Nb) was based on a modified version of the above protocol. Coverslips were placed on ice, washed with cold HKM-E1 buffer (20 mM HEPES, pH 7.4, 115 mM KOAc, 1 mM MgCl_2_, and 1 mM EGTA), and then incubated for 10 min in HKM-E1 buffer supplemented with 0.1% BSA and with 70 nM Alexa Fluor 647-labeled anti-GFP Nb (Breslow et al., 2013; Kirchhofer et al., 2010). After incubation with GFP Nb, coverslips were washed three times with HKM-E1 buffer, fixed in 4% paraformaldehyde, and shifted to room temperature. Coverslips were then permeabilized with PBS containing 0.1% Triton X-100 for 10 min, blocked, stained with anti-Rab34 antibody (Abcam), and processed as described above.

### Sample preparation for expansion microscopy

Cells seeded on 25mm #1.5 coverslips (Electron Microscopy Sciences) and prepared as in Sahabandu et al. (2019). Briefly, cells were fixed in 4% formaldehyde for 1 h, incubated in 30% acrylamide/4% formaldehyde at 37°C for 16 h, washed 3x 10min in PBS and then cooled on ice water. Pre-cooled gelling solution (20% Acrylamide, 0.04% bis-acrylamide, 7% sodium acrylate. 0.5% APS and 0.5% TEMED) was then added, and polymerization was allowed to proceed for 30 min on ice and 30-60 min at room temperature. 4-mm gel punches were taken using a biopsy punch (Integra Miltex, 33-34-P/25) and then denatured in SDS solution (50 mM Tris pH 9.0, 200 mM SDS, 200 mM NaCl) for 1 h at 90°C. Gel punches were allowed to cool, and SDS solution was washed out with PBS (9x 20 min washes) followed by incubation overnight in PBS at 4°C.

Denatured punches were then blocked in immunofluorescence buffer (1% BSA, 0.05% Tween-20 in PBS) for 2 h and incubated overnight at 4°C with primary antibodies in immunofluorescence buffer (see Supplementary Table 1). Punches were washed in PBS (6x 10 min), then incubated with secondary antibodies and Hoechst 33258 overnight at 4°C. Finally, stained punches were washed in water (12x 10 min), then expanded overnight in deionized water at 4°C. Gels were transferred to 35mm glass-bottom imaging dishes (Mattek P35G-1.5-14-C), immobilized using low-melt agarose, and imaged as described below. We estimate from the final gel size and the size of expanded nuclei that gels expanded ~4-fold, as previously reported (Sahabandu et al., 2019).

### Fluorescence microscopy

Coverslips with fixed NIH-3T3 and RPE cells were imaged using a Nikon Eclipse Ti-2 widefield microscope equipped with a CMOS camera (Photometrics Prime BSI or Hamamatsu Orca-Fusion), a 60✕ PlanApo oil objective (NA 1.40; Nikon Instruments), and an LED light source (Lumencor Spectra X or SOLA-V-NIR). Fixed samples were mounted in Fluoromount-G and images acquired at room temperature or 37°C using Nikon Elements software. MDCK cells cultured as 2D monolayers were imaged on an Olympus FV1000 confocal microscope using a 60✕ UPLSAPO oil objective (NA 1.35) controlled with Fluoview software (Olympus). MDCK cells cultured as 3D cysts were imaged on an Olympus IX83 microscope equipped with a Dragonfly200 spinning disk unit (Andor), a CMOS camera (Andor Zyla 4.2 Plus), a 60✕ UPLSAPO silicone oil objective (NA 1.30), and Fusion software (Andor). Fixed 2D MDCK cells were mounted in ProLong Diamond (Thermo Fisher) and images were acquired at room temperature.

Live-cell imaging and expansion microscopy imaging were performed on a Nikon Eclipse Ti-2 equipped a with Yokogawa W1 spinning disk unit, CMOS camera (Photometrics Prime BSI), 40✕ PlanApo silicone oil objective (NA 1.25), and 4-color laser combiner (Nikon Instruments LUN-F XL 405/488/561/640). For live-cell imaging, 42,000 cells per well were seeded on a μ-slide 8 well plate (Ibidi 80826), and after 24 h cells were imaged in phenol-red-free DMEM/F-12 medium with 10-1000 ng/ml doxycycline and 0.2% serum at 37°C using Nikon Elements software. A Perfect Focus System was used to maintain focus, and a stage-top incubator (Tokai Hit) was used to maintain temperature and 5% CO_2_. This system was used for all live-cell imaging, except for imaging of Ehd1-mScarlet, which was performed on the widefield Eclipse Ti-2 used for fixed-cell imaging (using an Okolab enclosure to maintain cells at 37°C and 5% CO_2_). For expansion microscopy, samples were imaged at room temperature and the supplemental 1.5x magnification tube lens of the Eclipse Ti-2 was used with the 40✕ PlanApo silicone oil objective (NA 1.25).

Structured illumination microscopy (SIM) imaging was performed on a Deltavision OMX v3 microscope (Applied Precision) equipped with a U-PLANAPO 60✕ oil objective (NA 1.42; Olympus), CoolSNAP HQ_2_ CCD cameras with a pixel size of 0.080 μm (Photometrics), and solid-state lasers at 488, 561, and 642 nm (Coherent and MPB communications). Samples were illuminated by a coherent scrambled laser light source that had passed through a diffraction grating to generate the structured illumination by interference of light orders in the image plane to create a 3D sinusoidal pattern, with lateral stripes approximately 270 nm apart. The pattern was shifted laterally through five phases and through three angular rotations of 60° for each z section (separated by 125 nm). Cells were stained with antibodies to Rab34 (Abcam) and ARL13B (NeuroMab), mounted in Prolong Diamond medium (Invitrogen P36965), and imaged at room temperature. Raw images were processed and deconvolved using Softworx software (Applied Precision). Channels acquired on separate cameras were then aligned in x, y, z, and rotationally using predetermined shifts as measured using the SoftWorx alignment tool. Tetraspeck beads (Invitrogen T7279) were used as a reference for channel alignment.

### Focused ion beam scanning electron microscopy (FIB-SEM)

Cells cultured on 12mm glass coverslips were fixed in 4% PFA for 20 min, washed in PBS, and further fixed in 2% glutaraldehyde in 0.1 M sodium cacodylate buffer, 2% OsO_4_ and 1.5% K_4_Fe(CN)_6_ (Sigma-Aldrich). Fixed cells were then stained en bloc in 2% aqueous uranyl acetate, dehydrated and embedded in Embed 812. Unless otherwise noted, all FIB-SEM reagents are from Electron Microscopy Sciences.

After removing the glass coverslip, the embedded monolayer of cultured cells was glued with EMS water-based conductive graphene carbon paint onto the sample mounting aluminum stub, with the coverslip-facing side of the cells exposed to the air. A 20-25 nm-thick conductive platinum coating was applied to the sample surface with a sputter coater (Ted Pella, Inc.), and the sample was then imaged in a Crossbeam 550 FIB-SEM workstation (Carl Zeiss Microscopy) operated with SmartSEM (Carl Zeiss Microscopy) or Atlas engine 5 (Fibics Incorporated, Ottawa, Canada) software. For image acquisition, a target area was first selected with the SE2 detector (Electron High Tension], 5kV; electron beam current, 2nA) at working distance of 10 nm, then the stage was inclined (to 54°) and lifted (to a working distance of 5 nm) as the eucentric height and coincident point were adjusted. The targeted cells’ surface (50×30 μm) was deposited with a 0.5-1 μm-thick protective platinum pad by using 30kV:3nA FIB. Autotune marks and 3D tracking marks were milled into the platinum pad, and carbon deposited by 30kV:50pA FIB, followed by deposition of another 0.5-1 μm-thick upper protective carbon pad. 30kV:30nA FIB was used for the coarse trench (20 μm deep), 30kV:3nA FIB was used for the fine trench, and 30kV:300pA FIB was used for milling the imaging surface during the 3D tracking. The dwell time for SEM imaging (Electron High Tension, 1.5 kV; electron beam current, 2 nA) with EsB detector was set at 2-3 μs with line averaging by 2. The filtering grid voltage of the EsB detector was set at 1000 V, and the collector voltage was set at 300 *V.* The imaging resolution was set at 7 nm/pixel at X, Y axis, slicing by 7 nm along Z axis, yielding final data sets with isotropic resolution at 7 nm/voxel. After the FIB-SEM imaging, the data sets were aligned and exported with Atlas 5 software, then further cropped and rotated with DragonFly software (Object Research Systems Inc.).

### Image analysis

Widefield fluorescence microscopy images were analyzed using Fiji/ImageJ. Ciliogenesis and quantification of protein markers present at cilia or centrioles was performed by manual counting. For SIM microscopy, images that were deconvolved using Softworx software (see above) were further aligned by imaging Tetraspeck beads and using the Fiji Descriptor-based registration (2D)’ plugin to determine and apply X/Y corrections. Intensity values for line traces across SIM images were determined in Fiji/Image using the ‘Plot Profile’ tool and plotted using Matlab (Mathworks, Inc.).

FIB-SEM datasets were processed using IMOD software (Kremer et al., 1996; Mastronarde and Held, 2017). Rotated sub-volumes were extracted using the ‘Rubber band’ tool. Centrioles and associated membranes were segmented manually using the ‘Sculpt’ drawing tool and rendered with the Surface and Cap Meshing options.

### Transfection, immunoprecipitation, SDS-PAGE, and Western blotting

HEK293T cells were transfected with LAP-Rab34 variants using X-tremeGENE 9 (Sigma Aldrich, 6365779001). After 48 h, the cells were collected, lysed on ice in Co-IP buffer (50 mM Tris pH 7.4, 150 mM NaCl, 1% Triton X-100, 0.5 mM DTT, protease inhibitors), and centrifuged at 20,000 x *g* for 15 min. LAP-tagged proteins were captured on GFP nanobody beads (Chromotek #gta-20). After washing, the beads were resuspended in 2X NuPAGE LDS buffer (Invitrogen) and denatured at 95°C for five min to elute captured proteins. For analysis of Rab34 levels in whole-cell lysate, cells were lysed in RIPA buffer (50 mM Tris pH 7.4, 150 mM NaCl, 2% NP40, 0.25% sodium deoxycholate, 0.5 mM DTT, protease inhibitors) on ice for 10 min and centrifuged at 20,000 x *g* for 10 min. The supernatant was collected, and equal amounts of protein were loaded, as determined by Bio-Rad Protein Assay.

Protein samples were separated on NuPAGE 4-12% Bis-Tris gels (Invitrogen) run in MOPS buffer (50 mM Tris, 50 mM MOPS, 3.5 mM SDS, 1 mM EDTA) and transferred to a PVDF membrane (Millipore Sigma #IPVH00010). The membranes were blocked in 1:1 PBS/SeaBlock (Thermo Fisher #37527) and incubated with primary antibodies overnight at 4°C (Supplementary Table 1). After washing, the membranes were incubated with HRP conjugated secondary antibodies or Protein A-HRP for 30 min at room temperature. The blots were developed using Clarity Western ECL Substrate (Bio-Rad) and imaged on a ChemiDoc Touch imager (Bio-Rad).

### Analysis of GTP/GDP binding in cells

HEK293T cells were transfected with LAP-Rab34 variants using X-tremeGENE 9. After 48 h, the cells were incubated with 0.25 mCi of ^32^P orthophosphate (Perkin Elmer, NEX053H002MC) for 4 h in DMEM containing dialyzed FBS (Gibco A3382001). The labeled cells were washed with ice-cold PBS and lysed on ice in Co-IP buffer supplemented with 10 mM MgCl_2._ The lysate was clarified by centrifugation at 20,000 x *g* for 15 min at 4°C, and LAP-tagged proteins were captured on GFP nanobody beads. After washing, bound nucleotides were dissociated by incubation in Elution buffer (0.2% SDS, 4 mM EDTA, 2 mM DTT, 1 mM GDP, 1 mM GTP) for 6 min at 68°C. The eluate was then spotted on a PEI-cellulose TLC plate (Sigma Z122882) and developed in a chamber equilibrated with 0.75 M KH_2_PO_4_. The dried plate was exposed to a phosphorimaging screen for 24 h, imaged on a Typhoon FLA 9500 (GE Healthcare), and quantified with ImageQuant software (GE Healthcare).

### Recombinant protein expression and purification

pGEX-Rab34 plasmids were transformed into BL21 DE3 competent cells expressing GroEL/ES (from pGro7 plasmid; gift from Yong Xiong, Yale University) and selected with 100 μg/mL ampicillin and 25 μg/mL chloramphenicol. Transformed cells were inoculated into a 100 mL culture of LB with ampicillin and chloramphenicol and grown at 37°C for 16 h. This culture was diluted to OD_600_=0.02-0.06 and expanded to 4L, followed by growth at 37°C until OD_600_=0.6. The cultures were then cooled in an ice water bath for 15 min. 2g/L arabinose was added to induce pGro7 expression and the cultures were transferred to an 18°C incubator. After 30 min, 200 μM IPTG was added to induce the expression of GST-Rab34. The cultures were grown at 18°C overnight; cultures were then collected by centrifugation at 3000 x *g* for 15 min at 4°C, and the pellet was flash frozen.

Cell pellets were suspended in lysis buffer (10 mM Tris pH 7.3, 1 mM DTT, 150 mM NaCl, 5 mM MgCl_2_, 0.001% Triton X-100, 1mM PMSF, 0.2 mM GDP, Roche cOmplete EDTA-free protease inhibitors) and lysed with a microfluidizer. Cell debris was removed through a high-speed (20,000 x *g*) clarification spin. The resulting supernatant was incubated with Glutathione Sepharose resin (GE Healthcare, 17-0756-05) for 2 h in lysis buffer. Bound Rab34 was eluted with elution buffer (10 mM Tris pH 8.2, 10 mM glutathione, 1 mM DTT, 70 mM NaCl, 5 mM MgCl_2_). Rab34 eluted from the affinity column was diluted 1:5 with Buffer A (10 mM Tris pH 8.2, 1 mM DTT, 70 mM NaCl, 5 mM MgCl_2_), loaded onto an anion exchange column (GE Healthcare HiTrap Q, 17-5156-01), washed with 10 column volumes of Buffer A, and eluted with a linear gradient of 0% to 100% Buffer B (10 mM Tris pH 8.2, 1 mM DTT, 0.5 M NaCl, 5 mM MgCl_2_) over 20 column volumes. Rab34 containing fractions were dialyzed overnight into storage buffer (50 mM Tris pH 7.5, 1 mM DTT, 150 mM NaCl, 5 mM MgCl_2_, 0.2 mM GDP) and simultaneously treated with HRV3C Protease (Pierce 88946) to cleave the GST tag. Dialyzed Rab34 was then concentrated to approximately 70 μM with an Amicon concentrator (Millipore, Z740199). Free GST tag, uncleaved GST-Rab34, and HRV3C were removed with a second round of incubation with Glutathione Sepharose resin. Lastly, glycerol was added to a final concentration of 40% (v/v) and the purified Rab34 was stored at −20°C. All steps were conducted at 4°C unless noted otherwise.

### Analysis of GTPase activity in vitro

All Rab34 used in steady-state assays was treated with AG 1-X8 anion exchange resin (Bio-Rad, 1401441) immediately before use to remove bound GDP. Steady-state GTPase activity was measured with a modified version of the standard NADH-based assay (Bradley and De La Cruz, 2012; Ingerman and Nunnari, 2005) in which the final concentrations of coupling reaction components were adjusted as follows: Pyruvate kinase (Sigma Aldrich, P9136) – 400 units/mL, Lactate dehydrogenase (BBI Enzymes, LDHP2FS) – 40 units/mL, Phosphoenolpyruvate (Sigma Aldrich, 10108294) – 10 mM. Rab34 GTPase activity was assayed in Rab34 assay buffer (50 mM Tris pH 7.7, 150 mM NaCl, 5 mM MgCl_2_, 1 mM DTT) at 25°C. Absorbance at 340 nm was measured on an Olis HP 8452 Diode Array spectrophotometer.

### Statistical analysis

Statistical tests were carried out using t-test (two-sided, paired) or Tukey-Kramer test, as indicated in Figure Legends.

## Supplemental material

Supplementary Figure 1 shows additional characterization of *RAB34* knockout cells. Supplementary Figure 2 shows conservation of atypical sequence elements in Rab34, biochemical analysis of Rab34 GTPase, and further characterization of the ciliogenesis defect observed in cells expressing Rab34-S166N. Supplementary Figure 3 presents additional analysis of Rab34 localization to intracellular versus externally exposed cilia, as assessed by the In/Out assay. Additional 3D-SIM and expansion microscopy imaging of Rab34 is also included.

Supplementary Table 1 lists antibodies used, and Supplementary Table 2 lists oligonucleotide and sgRNA sequences.

## Acknowledgments

We acknowledge David Mick and members of the D.K.B. lab for advice and discussion; Derek Toomre (Yale University) for In/Out reporter construct and cell line; Chris Westlake (National Cancer Institute) for cDNA encoding Ehd1; Tamara Caspary (Emory University) for Arl13b cDNA; Tim Stearns (Stanford University) for Centrin2 cDNA; and Yong Xiong (Yale University) for pGro7 plasmid. This work was supported by funding from the National Institutes of Health (R00HD082280 and R35GM137956 to D.K.B.; R35GM136656 to E.M.D.L.C.), the Charles H. Hood Foundation (to D.K.B.), the Alfred P. Sloan Foundation (FG-2018-10333 to D.K.B.), an Anderson Endowed Fellowship (A.K.G.), the Ministry of Education, Culture, Sports, Science and Technology (MEXT) of Japan (Grant-in-Aid for Young Scientists 20K15739 to Y. H.; Grant-in-Aid for Scientific Research(B) 19H03220 to M. F.), the Japan Science and Technology Agency (CREST grant JPMJCR17H4 to M. F.), the Japan Society for the Promotion of Science (to M. E. O), and NSF major instrumentation grant 1725480. S.D.G. was supported by R35GM136656-S1. We acknowledge Felix Rivera-Molina, Derek Toomre, and the Yale CINEMA microscopy facility for assistance with SIM imaging; Yumei Wu, Xinran Liu, and the Yale Center for Cellular and Molecular Imaging for FIB-SEM sample processing and data collection; Ken Nelson and the MCDB FACS facility for assistance with cell sorting; Shirin Bahmanyar, Jing Yan, and Nadya Dimitrova for microscope use; and Anna Pyle for phosphorimager use. The authors declare no competing financial interests.

## Author Contributions

Anil Kumar Ganga and Margaret C. Kennedy designed and conducted most experiments and analyzed results. Mai E. Oguchi, Yuta Homma, and Mitsunori Fukuda analyzed ciliogenesis in MDCK cells. Kendall Oliver analyzed Rab34 mutants, and Tracy Knight conducted expansion microscopy. Shawn D. Gray developed the recombinant protein purification protocol and performed biochemical analyses with purified protein components under the supervision of Enrique M. De La Cruz. David Breslow designed experiments, analyzed results, and supervised the project. David K. Breslow and Anil Kumar Ganga wrote the manuscript with input from all authors.

## Abbreviations

cytoD: cytochalasin D
DAV: Distal appendage vesicle
FIB-SEM: Focused ion beam scanning electron microscopy
GAP: GTPase activating protein
GEF: Guanine nucleotide exchange factor
Hh: Hedgehog
LAP: Localization and affinity purification tag (containing GFP, TEV protease site, and S-tag)
Nb: Nanobody
GEF: Guanine nucleotide exchange factor
polyE-tub: polyglutamylated tubulin
sgRNA: single guide RNA
SIM: Structured illumination microscopy

---

**Supplementary Figure 1.**
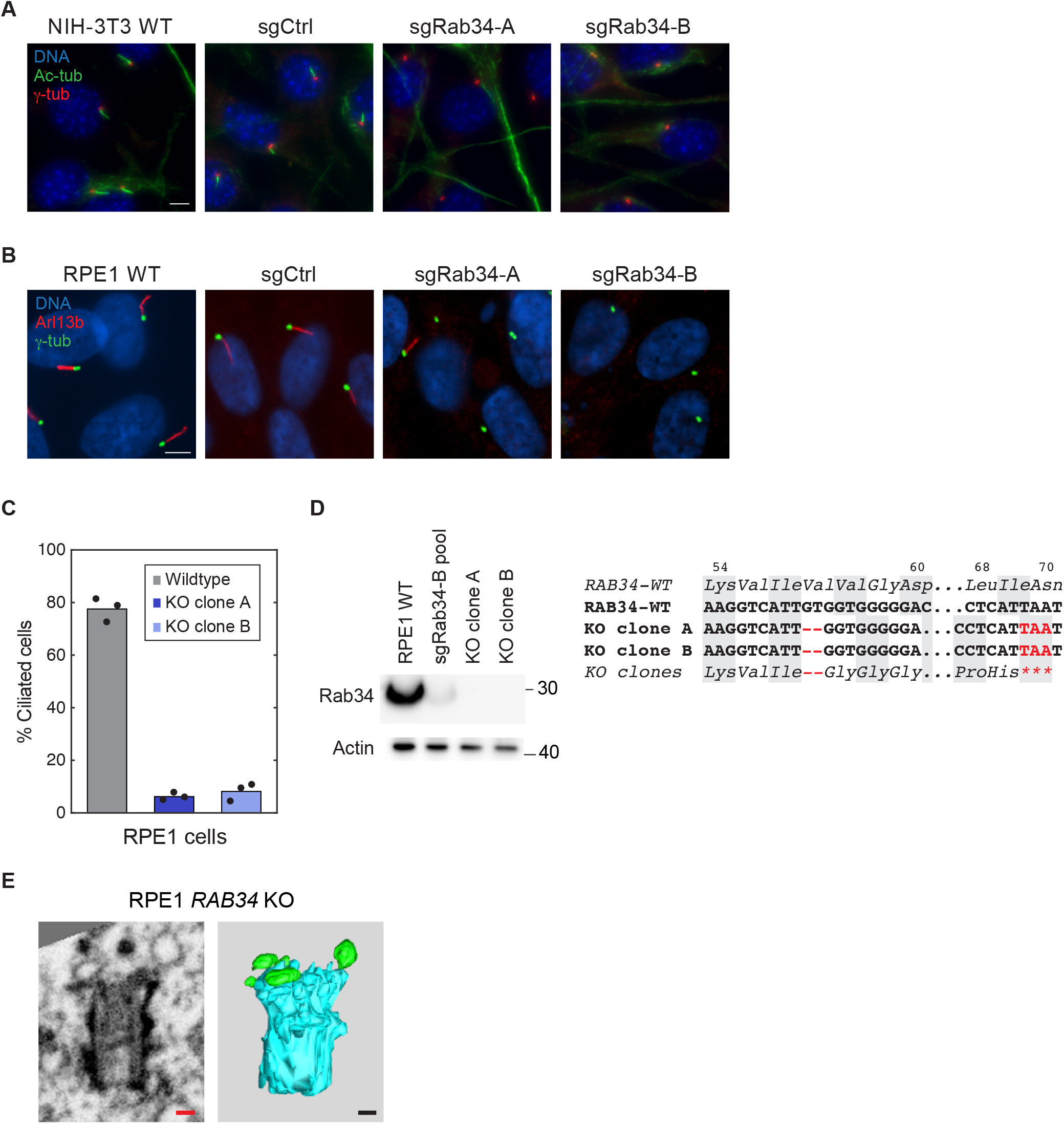
Characterization of Rab34 knockout cells and sequence conservation. **A**) Representative images of cilia (marked by acetylated tubulin; Ac-tub) and centrioles in NIH-3T3 cells transduced with the indicated sgRNAs. Scale bars: 5 μm. **B**) Representative images of cilia and centrioles in RPE1 cells transduced with the indicated sgRNAs. Scale bars: 5 μm. **C**) Quantification of ciliogenesis in WT RPE1 cells and clonal knockout cells obtained following transduction with sgRab34-2. Ciliation was assessed by ARL13B and γ-tubulin staining. Dots indicate values from individual experiments and bars indicate mean from *N*=3 independent experiments. **D**) Western blot of Rab34 in RPE1 WT cells and the indicated Rab34 mutants (left). DNA sequence of WT *RAB34* locus and of homozygous mutations observed in RPE1 Rab34 knockout clones (right). **E**) Additional representative FIB-SEM image and 3D reconstruction obtained from RPE1 *RAB34* knockout cells. Scale bars indicate 100 nm. See also Figure 2 G.

**Supplementary Figure 2.**
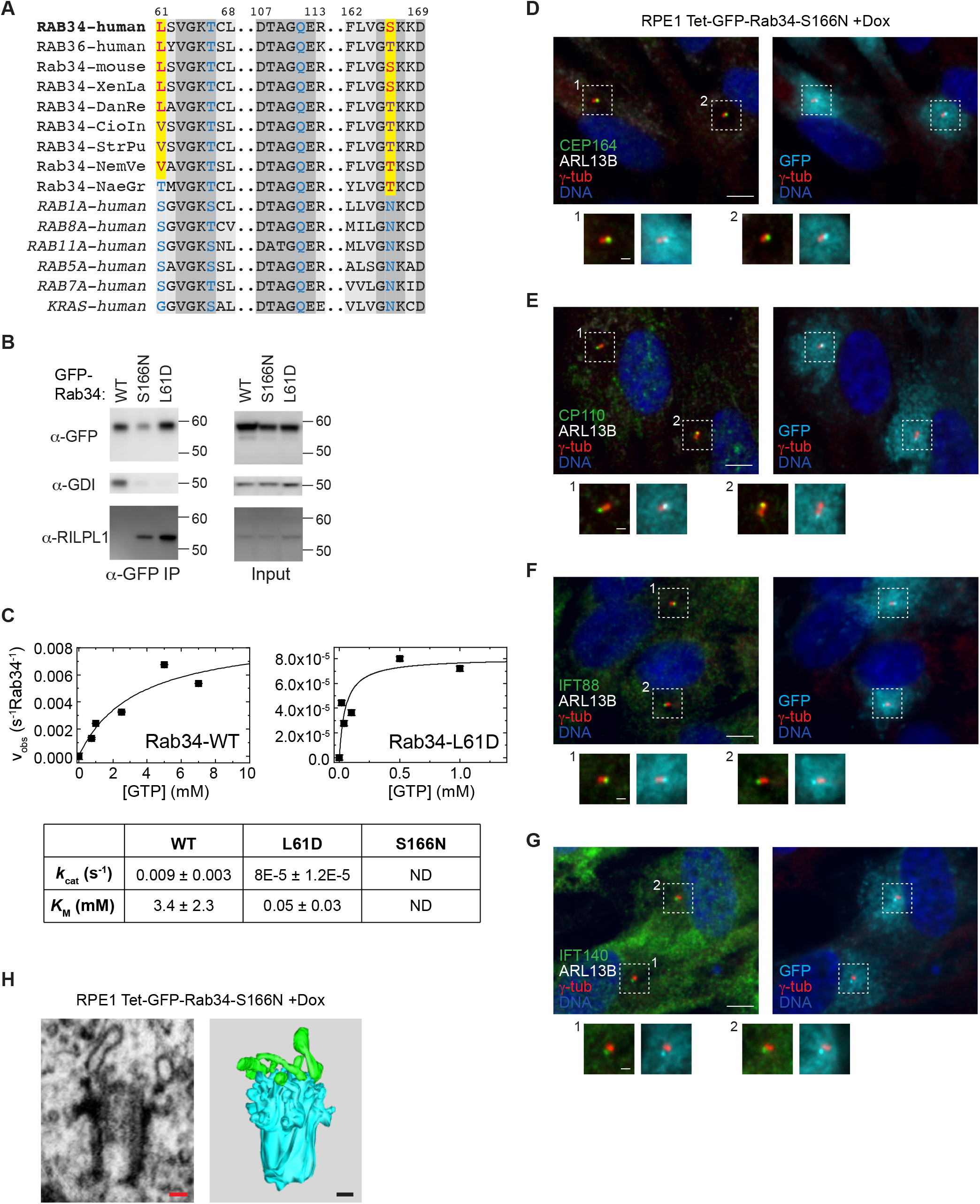
Characterization of the ciliogenesis defect induced by Rab34-S166N. **A**) Sequence comparison of human Rab34 to its close paralog Rab36 and to homologs from the indicated species. Residues conserved in Rab34 family members that diverge from canonical sequence are highlighted yellow while the canonical amino acid is shown in blue. Abbreviations: XenLa = *Xenopus laevis*; DanRe = *Danio rerio*; CioIn = *Ciona Intestinalis* (sea squirt); StrPu = *Strongylocentrotus purpuratus* (sea urchin); NemVe = *Nematostella vectensis* (sea anemone); NaeGr = *Naegleria gruberi*. **B**) HEK-293T cells were transfected with the indicated GFP-Rab34 variants and the indicated proteins were detected by Western blot in whole cell lysate (Input) or following anti-GFP immunoprecipitation (α-GFP IP). One of three representative experiments. **C**) Steady-state GTP hydrolysis is shown for purified Rab34-WT and Rab34-L61D (top). Uncertainty bars represent standard error of the fits to time-dependent NADH absorption readings. The continuous line through the data represents the best fit to a rectangular hyperbola, yielding the maximum velocity per enzyme (k_cat_) and the K_M,GTP_. Uncertainties in k_cat_ and K_M,GTP_ represent standard error in the fits to the rectangular hyperbola. A summary of results is shown (below); no measurable hydrolysis was observed for Rab34-S166N, and k_cat_ and K_M,GTP_ are reported as not detected (ND). One of *N*=3 representative experiments. **D-G**) Localization of CEP164, CP110, IFT88, and IFT140 was assessed in RPE Tet-GFP-Rab34 cells following 48 h serum starvation in the presence of doxycycline. Scale bars: 5 μm (insets: 1μm). **H**) Additional example of FIB-SEM analysis of centrioles and associated membranes in RPE1 GFP-Rab34-S166N cells. See also Figure 3 F.

**Supplementary Figure 3.**
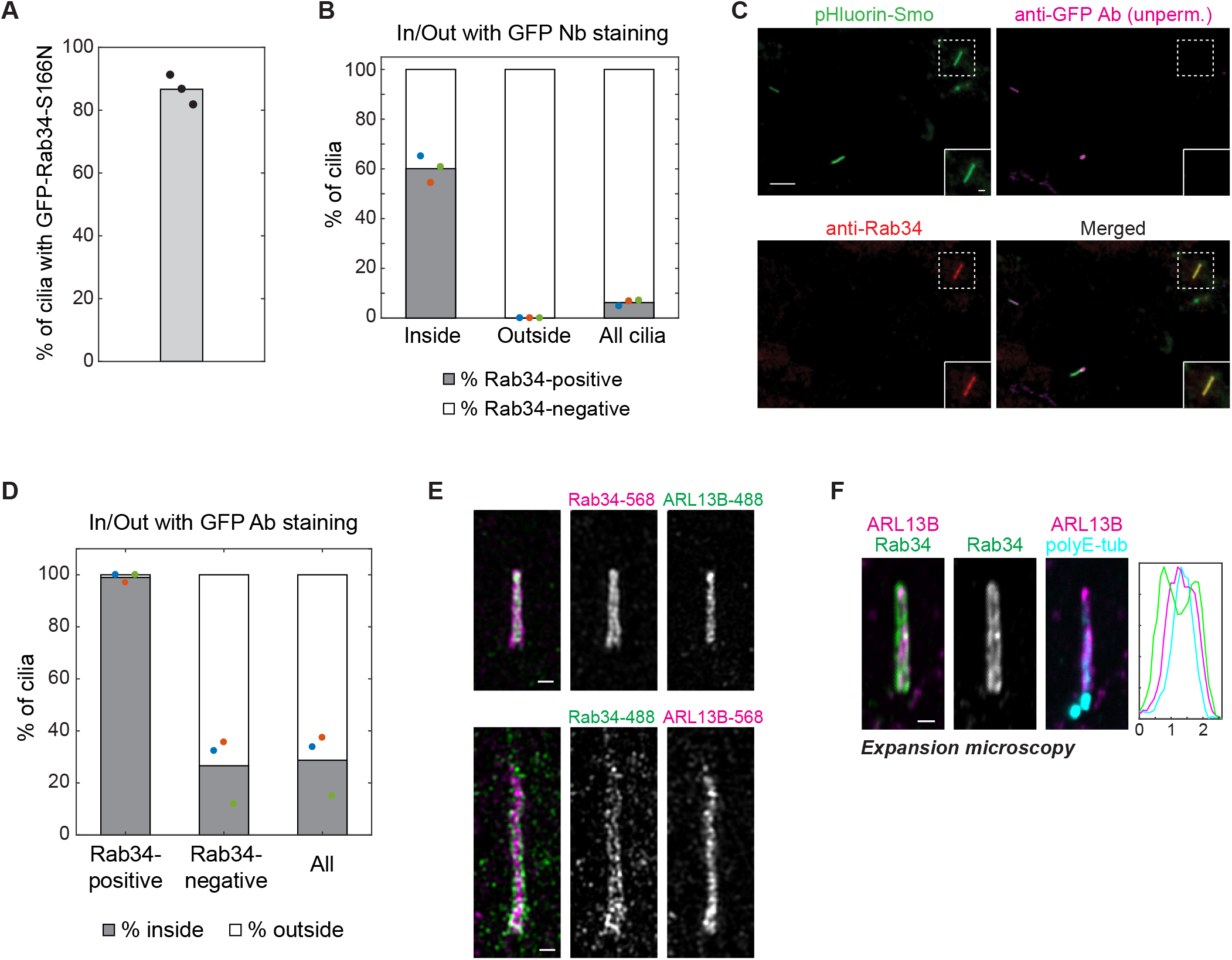
Rab34 localizes to the ciliary sheath of intracellular cilia. **A)** Quantification of RPE1 Tet-GFP-Rab34-S166N cells exhibiting GFP localization to the mother centriole after doxycycline induction and 48 h serum starvation. *N*=3 independent experiments. **B)** Quantification of the fraction of cilia that are Rab34-positive versus Rab34-negative is shown for the cilia classes indicated. Data shown are from the same GFP nanobody-based In/Out assay shown in Figure 5 A-B. Dots indicate values from individual experiments and bars indicate mean from *N*=3 independent experiments. **C**) RPE1 cells expressing the pHluorin-Smo In/Out reporter were stained prior to permeabilization with anti-GFP antibody (Ab) to reveal intracellular versus extracellular cilia followed by permeabilization and staining for Rab34 (SCBT antibody). Inset shows an intracellular (inside) cilium that is positive for Rab34. Scale bars: 5 μm (insets: 1μm). **D**) Quantification of the fraction of cilia that are inside versus outside is shown for the cilia classes indicated, as assessed using the In/Out assay shown in (**C**). **E**) Additional representative cilia are shown for the analysis presented in Figure 5 C. 3D-SIM (structured illumination microscopy) was performed on RPE1 cells stained for Rab34 and ARL13B. Alexa Fluor 488- and Alexa Fluor 568-conjugated secondary antibodies were used for both target proteins to ensure consistent results independent of emission wavelength. Graphs at right show normalized fluorescence intensity along line-profiles perpendicular to the longitudinal axis of the cilium. Scale bars indicate 100 nm. **F**) Expansion microscopy staining as in Figure 3 D with polyE-tubulin included. Scale bar indicates 2μm (after ~4-fold expansion).

## References

Anvarian, Z., K. Mykytyn, S. Mukhopadhyay, L.B. Pedersen, and S.T. Christensen. 2019. Cellular signalling by primary cilia in development, organ function and disease. Nat Rev Nephrol. 15:199–219.

Bangs, F., and K.V. Anderson. 2017. Primary Cilia and Mammalian Hedgehog Signaling. Cold Spring Harb Perspect Biol. 9:a028175.

Bernabe-Rubio, M., and M.A. Alonso. 2017. Routes and machinery of primary cilium biogenesis. Cell Mol Life Sci. 74:4077–4095.

Bernabe-Rubio, M., G. Andres, J. Casares-Arias, J. Fernandez-Barrera, L. Rangel, N. Reglero-Real, D.C. Gershlick, J.J. Fernandez, J. Millan, I. Correas, et al. 2016. Novel role for the midbody in primary ciliogenesis by polarized epithelial cells. J Cell Biol. 214:259–273.

Blacque, O.E., N. Scheidel, and S. Kuhns. 2018. Rab GTPases in cilium formation and function. Small GTPases. 9:76–94.

Bradley, M.J., and E.M. De La Cruz. 2012. Analyzing ATP utilization by DEAD-Box RNA helicases using kinetic and equilibrium methods. Methods Enzymol. 511:29–63.

Braun, D.A., and F. Hildebrandt. 2017. Ciliopathies. Cold Spring Harb Perspect Biol. 9:a028191.

Breslow, D.K., and A.J. Holland. 2019. Mechanism and Regulation of Centriole and Cilium Biogenesis. Annu Rev Biochem. 88:691–724.

Breslow, D.K., S. Hoogendoorn, A.R. Kopp, D.W. Morgens, B.K. Vu, M.C. Kennedy, K. Han, A. Li, G.T. Hess, M.C. Bassik, et al. 2018. A CRISPR-based screen for Hedgehog signaling provides insights into ciliary function and ciliopathies. Nat Genet. 50:460–471.

Breslow, D.K., E.F. Koslover, F. Seydel, A.J. Spakowitz, and M.V. Nachury. 2013. An in vitro assay for entry into cilia reveals unique properties of the soluble diffusion barrier. J Cell Biol. 203:129–147.

Chang, C.H., M. Zanini, H. Shirvani, J.S. Cheng, H. Yu, C.H. Feng, A.L. Mercier, S.Y. Hung, A. Forget, C.H. Wang, et al. 2019. Atoh1 Controls Primary Cilia Formation to Allow for SHH-Triggered Granule Neuron Progenitor Proliferation. Dev Cell. 48:184–199 e185.

Dickinson, M.E., A.M. Flenniken, X. Ji, L. Teboul, M.D. Wong, J.K. White, T.F. Meehan, W.J. Weninger, H. Westerberg, H. Adissu, et al. 2016. High-throughput discovery of novel developmental phenotypes. Nature. 537:508–514.

Diekmann, Y., E. Seixas, M. Gouw, F. Tavares-Cadete, M.C. Seabra, and J.B. Pereira-Leal. 2011. Thousands of rab GTPases for the cell biologist. PLoS Comput Biol. 7:e1002217.

Fukuda, M., E. Kanno, K. Ishibashi, and T. Itoh. 2008. Large scale screening for novel rab effectors reveals unexpected broad Rab binding specificity. Mol Cell Proteomics. 7:1031–1042.

Garcia, G., 3rd, D.R. Raleigh, and J.F. Reiter. 2018. How the Ciliary Membrane Is Organized Inside-Out to Communicate Outside-In. Curr Biol. 28:R421–R434.

Gerondopoulos, A., H. Strutt, N.L. Stevenson, T. Sobajima, T.P. Levine, D.J. Stephens, D. Strutt, and F.A. Barr. 2019. Planar Cell Polarity Effector Proteins Inturned and Fuzzy Form a Rab23 GEF Complex. Curr Biol. 29:3323–3330 e3328.

Ghossoub, R., A. Molla-Herman, P. Bastin, and A. Benmerah. 2011. The ciliary pocket: a once-forgotten membrane domain at the base of cilia. Biol Cell. 103:131–144.

Haycraft, C.J., B. Banizs, Y. Aydin-Son, Q. Zhang, E.J. Michaud, and B.K. Yoder. 2005. Gli2 and Gli3 localize to cilia and require the intraflagellar transport protein polaris for processing and function. PLoS Genet. 1:e53.

Homma, Y., R. Kinoshita, Y. Kuchitsu, P.S. Wawro, S. Marubashi, M.E. Oguchi, M. Ishida, N. Fujita, and M. Fukuda. 2019. Comprehensive knockout analysis of the Rab family GTPases in epithelial cells. J Cell Biol. 218:2035–2050.

Huangfu, D., A. Liu, A.S. Rakeman, N.S. Murcia, L. Niswander, and K.V. Anderson. 2003. Hedgehog signalling in the mouse requires intraflagellar transport proteins. Nature. 426:83–87.

Ingerman, E., and J. Nunnari. 2005. A continuous, regenerative coupled GTPase assay for dynamin-related proteins. Methods Enzymol. 404:611–619.

Insinna, C., Q. Lu, I. Teixeira, A. Harned, E.M. Semler, J. Stauffer, V. Magidson, A. Tiwari, A.K. Kenworthy, K. Narayan, et al. 2019. Investigation of F-BAR domain PACSIN proteins uncovers membrane tubulation function in cilia assembly and transport. Nat Commun. 10:428.

Kirchhofer, A., J. Helma, K. Schmidthals, C. Frauer, S. Cui, A. Karcher, M. Pellis, S. Muyldermans, C.S. Casas-Delucchi, M.C. Cardoso, et al. 2010. Modulation of protein properties in living cells using nanobodies. Nat Struct Mol Biol. 17:133–138.

Knodler, A., S. Feng, J. Zhang, X. Zhang, A. Das, J. Peranen, and W. Guo. 2010. Coordination of Rab8 and Rab11 in primary ciliogenesis. Proc Natl Acad Sci U S A. 107:6346–6351.

Kremer, J.R., D.N. Mastronarde, and J.R. McIntosh. 1996. Computer visualization of three-dimensional image data using IMOD. J Struct Biol. 116:71–76.

Kukic, I., F. Rivera-Molina, and D. Toomre. 2016. The IN/OUT assay: a new tool to study ciliogenesis. Cilia. 5:23.

Langemeyer, L., R. Nunes Bastos, Y. Cai, A. Itzen, K.M. Reinisch, and F.A. Barr. 2014. Diversity and plasticity in Rab GTPase nucleotide release mechanism has consequences for Rab activation and inactivation. Elife. 3:e01623.

Leaf, A., and M. Von Zastrow. 2015. Dopamine receptors reveal an essential role of IFT-B, KIF17, and Rab23 in delivering specific receptors to primary cilia. Elife. 4:e06996.

Lee, E.Y., H. Ji, Z. Ouyang, B. Zhou, W. Ma, S.A. Vokes, A.P. McMahon, W.H. Wong, and M.P. Scott. 2010. Hedgehog pathway-regulated gene networks in cerebellum development and tumorigenesis. Proc Natl Acad Sci U S A. 107:9736–9741.

Lee, M.T., A. Mishra, and D.G. Lambright. 2009. Structural mechanisms for regulation of membrane traffic by rab GTPases. Traffic. 10:1377–1389.

Lu, Q., C. Insinna, C. Ott, J. Stauffer, P.A. Pintado, J. Rahajeng, U. Baxa, V. Walia, A. Cuenca, Y.S. Hwang, et al. 2015. Early steps in primary cilium assembly require EHD1/EHD3-dependent ciliary vesicle formation. Nat Cell Biol. 17:228–240.

Mastronarde, D.N., and S.R. Held. 2017. Automated tilt series alignment and tomographic reconstruction in IMOD. J Struct Biol. 197:102–113.

Matsui, T., N. Ohbayashi, and M. Fukuda. 2012. The Rab interacting lysosomal protein (RILP) homology domain functions as a novel effector domain for small GTPase Rab36: Rab36 regulates retrograde melanosome transport in melanocytes. J Biol Chem. 287:28619–28631.

Mirvis, M., T. Stearns, and W. James Nelson. 2018. Cilium structure, assembly, and disassembly regulated by the cytoskeleton. Biochem J. 475:2329–2353.

Mizuno-Yamasaki, E., F. Rivera-Molina, and P. Novick. 2012. GTPase networks in membrane traffic. Annu Rev Biochem. 81:637–659.

Muller, M.P., and R.S. Goody. 2018. Molecular control of Rab activity by GEFs, GAPs and GDI. Small GTPases. 9:5–21.

Nachury, M.V., A.V. Loktev, Q. Zhang, C.J. Westlake, J. Peranen, A. Merdes, D.C. Slusarski, R.H. Scheller, J.F. Bazan, V.C. Sheffield, et al. 2007. A core complex of BBS proteins cooperates with the GTPase Rab8 to promote ciliary membrane biogenesis. Cell. 129:1201–1213.

Nachury, M.V., and D.U. Mick. 2019. Establishing and regulating the composition of cilia for signal transduction. Nat Rev Mol Cell Biol. 20:389–405.

Nottingham, R.M., and S.R. Pfeffer. 2014. Mutant enzymes challenge all assumptions. Elife. 3:e02171.

Oguchi, M.E., K. Okuyama, Y. Homma, and M. Fukuda. 2020. A comprehensive analysis of Rab GTPases reveals a role for Rab34 in serum starvation-induced primary ciliogenesis. J Biol Chem. 295:12674–12685.

Pfeffer, S.R. 2017. Rab GTPases: master regulators that establish the secretory and endocytic pathways. Mol Biol Cell. 28:712–715.

Prior, I.A., P.D. Lewis, and C. Mattos. 2012. A comprehensive survey of Ras mutations in cancer. Cancer Res. 72:2457–2467.

Pusapati, G.V., J.H. Kong, B.B. Patel, A. Krishnan, A. Sagner, M. Kinnebrew, J. Briscoe, L. Aravind, and R. Rohatgi. 2018. CRISPR Screens Uncover Genes that Regulate Target Cell Sensitivity to the Morphogen Sonic Hedgehog. Dev Cell. 44:113–129 e118.

Reiter, J.F., and M.R. Leroux. 2017. Genes and molecular pathways underpinning ciliopathies. Nat Rev Mol Cell Biol. 18:533–547.

Sahabandu, N., D. Kong, V. Magidson, R. Nanjundappa, C. Sullenberger, M.R. Mahjoub, and J. Loncarek. 2019. Expansion microscopy for the analysis of centrioles and cilia. J Microsc. 276:145–159.

Schaub, J.R., and T. Stearns. 2013. The Rilp-like proteins Rilpl1 and Rilpl2 regulate ciliary membrane content. Mol Biol Cell. 24:453–464.

Schmidt, K.N., S. Kuhns, A. Neuner, B. Hub, H. Zentgraf, and G. Pereira. 2012. Cep164 mediates vesicular docking to the mother centriole during early steps of ciliogenesis. J Cell Biol. 199:1083–1101.

Sorokin, S. 1962. Centrioles and the formation of rudimentary cilia by fibroblasts and smooth muscle cells. J Cell Biol. 15:363–377.

Sorokin, S.P. 1968. Reconstructions of centriole formation and ciliogenesis in mammalian lungs. J Cell Sci. 3:207–230.

Stenmark, H. 2009. Rab GTPases as coordinators of vesicle traffic. Nat Rev Mol Cell Biol. 10:513–525.

Sun, P., H. Yamamoto, S. Suetsugu, H. Miki, T. Takenawa, and T. Endo. 2003. Small GTPase Rah/Rab34 is associated with membrane ruffles and macropinosomes and promotes macropinosome formation. J Biol Chem. 278:4063–4071.

Tillberg, P.W., and F. Chen. 2019. Expansion Microscopy: Scalable and Convenient Super-Resolution Microscopy. Annu Rev Cell Dev Biol. 35:683–701.

Tisdale, E.J., J.R. Bourne, R. Khosravi-Far, C.J. Der, and W.E. Balch. 1992. GTP-binding mutants of rab1 and rab2 are potent inhibitors of vesicular transport from the endoplasmic reticulum to the Golgi complex. J Cell Biol. 119:749–761.

Vokes, S.A., H. Ji, S. McCuine, T. Tenzen, S. Giles, S. Zhong, W.J. Longabaugh, E.H. Davidson, W.H. Wong, and A.P. McMahon. 2007. Genomic characterization of Gli-activator targets in sonic hedgehog-mediated neural patterning. Development. 134:1977–1989.

Walworth, N.C., B. Goud, A.K. Kabcenell, and P.J. Novick. 1989. Mutational analysis of SEC4 suggests a cyclical mechanism for the regulation of vesicular traffic. EMBO J. 8:1685–1693.

Wandinger-Ness, A., and M. Zerial. 2014. Rab proteins and the compartmentalization of the endosomal system. Cold Spring Harb Perspect Biol. 6:a022616.

Wang, L., and B.D. Dynlacht. 2018. The regulation of cilium assembly and disassembly in development and disease. Development. 145:dev151407.

Wu, C.T., H.Y. Chen, and T.K. Tang. 2018. Myosin-Va is required for preciliary vesicle transportation to the mother centriole during ciliogenesis. Nat Cell Biol. 20:175–185.

Xu, S., Y. Liu, Q. Meng, and B. Wang. 2018. Rab34 small GTPase is required for Hedgehog signaling and an early step of ciliary vesicle formation in mouse. J Cell Sci. 131:jcs213710.

Ye, F., D.K. Breslow, E.F. Koslover, A.J. Spakowitz, W.J. Nelson, and M.V. Nachury. 2013. Single molecule imaging reveals a major role for diffusion in the exploration of ciliary space by signaling receptors. Elife. 2:e00654.

